# Widespread allele-specific topological domains in the human genome are not confined to imprinted gene clusters

**DOI:** 10.1101/2022.04.26.489502

**Authors:** Stephen Richer, Tian Yuan, Stefan Schoenfelder, Laurence Hurst, Adele Murrell, Giuseppina Pisignano

**Affiliations:** Dept Biology and Biochemistry, University of Bath, Claverton Down, Bath, BA2 7AY, United Kingdom; Babraham Institute, Cambridge, CB22 3AT, United Kingdom

## Abstract

**Background:** There is widespread interest in the three-dimensional chromatin conformation of the genome and its impact on gene expression. However, these studies frequently do not consider parent-of-origin differences, such as genomic imprinting, which result in monoallelic expression. In addition, genome-wide allele-specific chromatin conformation associations have not been extensively explored. There are few accessible bioinformatic workflows for investigating allelic conformation differences and these require pre-phased haplotypes which are not widely available.

**Results:** We assembled a bioinformatic pipeline, “HiCFlow”, which performs haplotype assembly and visualisation of parental chromatin architecture. We benchmarked the pipeline using prototype haplotype phased Hi-C data from GM12878 cells at three disease associated imprinted gene clusters. Using RC-HiC (Region Capture Hi-C) and Hi-C data from further human cell lines (1-7HB2, IMR-90, and H1-hESCs) we were able to robustly identify the known stable allele-specific interactions at the *H19/IGF2* locus. Other imprinted loci (*DLK1* and *SNRPN*) were more variable and there was no “canonical imprinted 3D structure”, but we could detect allele-specific differences in A/B compartmentalisation. Genome-wide, when TADs were unbiasedly ranked according to their allele-specific contact frequencies, a set of “allele-specific TADs” (ASTADs) could be defined. These occurred in genomic regions of high sequence variation. In addition to imprinted genes, ASTADs were also enriched for allele-specific expressed (ASE) genes. We found loci in ASTADs that have not previously been identified as ASE such as the bitter taste receptors (*TAS2R*s).

**Conclusions:** This study highlights the widespread differences in chromatin conformation between heterozygous loci and provides a new framework for understanding ASE.

## Background

Higher order chromatin conformation forms a scaffold (Dixon *et al*., 2012; Dorsett and Merkenschlager, 2013) upon which epigenetic mechanisms converge to regulate allele-specific gene expression. While chromatin capture approaches provide a snapshot of a high number of dynamic interactions, these are often averaged across both alleles in a heterogeneous cell population. Such approaches mask any allele-specific differences and may bias the interpretation of chromatin conformation at sites of allele-specific gene expression (ASE) (Buckland, 2004). However, ASE is known to occur widely in the human genome and is an important contributor to heritable differences in phenotypic traits. Monoallelic expression is a special case of ASE, regulated by an array of epigenetic mechanisms rather than genetic sequence of the allele (Reinius and Sandberg, 2015). Genomic imprinting is a well-studied example of monoallelic expression, and unique in that the active and silent alleles acquire their epigenetic marks during gametic development resulting in expression linked to the parental origin of the gene. “Silent” imprinted alleles adopt permanently repressed epigenetic signatures, whereas their “active” homologues can be expressed tissue specifically. Epigenetic elements of imprinted gene regulation include sequences that are methylated on only one of the parental alleles, (known as differentially methylated regions, DMRs). DMRs further have underlying allelic differences in post-translational histone modifications and ‘CCCTC-binding factor’ (CTCF) occupancy (reviewed by Monk *et al*., 2019).

Disturbances of the allelic dosage due to chromosomal rearrangements or the epigenetic disruption of co-regulated expression in imprinted genes, lead to defined clinical syndromes collectively known as imprinting disorders (reviewed by Monk *et al.,* 2019). The most common imprinting disorders include Beckwith Wiedemann syndrome (BWS) (Soejima and Higashimoto, 2013; Azzi, Habib and Netchine, 2014) with an incidence of 1 in 15,000 live births, Angelman Syndrome (AS, 1:20,000); Prader Willi Syndrome (PWS, 1:25,000) (Rabinovitz *et al*., 2012), and Silver Russell Syndrome (SRS, 1:100,000) (Azzi, Habib and Netchine, 2014).

The *IGF2-KCNQ1* locus, implicated in BWS and SRS, divides into two imprinted gene clusters. The first is *IGF2-H19*, which is regulated by an imprinting control region (ICR) known as the *H19*-DMR. The *H19*-DMR is methylated over CTCF binding sites on the paternal allele, thus preventing the paternal allele from forming CTCF mediated loops with this site. The second is the *KCNQ1OT1* cluster that is regulated by a DMR at the promoter of the *KCNQ1OT1* gene which encodes a long non-coding RNA (lncRNA) (Naveh *et al*., 2021). Transcription of this lncRNA on the unmethylated paternal allele silences *KCNQ1* and adjacent genes including *TSSC4, CDKN1* and *PHLDA2* (Korostowski, Sedlak and Engel, 2012). The *SNRPN* locus is implicated in Prader-Willi and Angelman syndromes. Imprinting at this region is regulated by a ∼35kb bipartite imprinting control region (PWS-IC and AS-IC). PWS-IC is the promoter for the pre-mRNA transcript for *SNRPN* and *SNURF* and the ncRNA *SNHG14*, which is a host transcript for several long and short non-coding RNAs such as *SPA1*, *SPA2*, sno-lncRNAs 1–5, *SNORD116*, *IPW*, *SNORD115* and *UBE3A-ATS*. PWS-IC is a DMR that overlaps the first exon of *SNURF-SNRPN* and is methylated on the maternal allele. The AS-IC is not a DMR but needs to be present to ensure the correct methylation at the PWS-IC (Horsthemke and Wagstaff, 2008). The *DLK1-DIO3* locus harbours *DLK1* and *RTL1 (*protein-coding paternally expressed genes) as well as several non-coding maternally expressed genes namely *MEG3* (alias, *GTL2*), *RTL1-AS*, *MEG8*, tandemly repeated array of C/D-box snoRNAs (small nucleolar RNAs), and more than 50 miRNAs (Benetatos *et al*., 2013). Between the *DLK1* gene and the transcription start site of the ncRNA cluster are two DMRs that regulate monoallelic expression: one is in the intergenic region between *DLK1* and *MEG3* (IG-DMR and the second is located ∼20kb away at the *MEG3* promoter (*MEG3*-DMR) (Kagami *et al*., 2010). These DMRs are methylated on the paternal allele in most somatic tissues. In humans, imprinting abnormalities and dosage effects at this imprinted region cause Temple (MatUPD14) and Kagami-Ogata (PatUPD14) syndromes (Ogata and Kagami, 2016; Prasasya *et al*., 2020) with phenotypes of postnatal growth retardation, skeletal malformations, and metabolic deficiencies (Temple *et al*., 1991; Kagami *et al*., 2010).

Imprinted genes are an excellent model system for analysing epigenetic regulation of gene expression and the study of genomic imprinting has uncovered many paradigms that are generally relevant to gene expression (Adalsteinsson and Ferguson-Smith, 2014). One such paradigm is that the CTCF can act as a boundary element separating different regulatory elements that could be shared between genes. We and others have shown that differential binding of CTCF-cohesin complexes at the imprinted *IGF2-H19* locus regulates access to a series of enhancers through allele-specific differences in higher order looping interactions (Bell and Felsenfeld, 2000; Hark *et al*., 2000; Szabó *et al.,* 2000; Kurukuti *et al.,* 2006; Engel *et al.,* 2008; Nativio *et al.,* 2009).

These early studies used a chromosome conformation capture (3C) technique in which fixed chromatin is digested with a restriction enzyme followed by a ligation reaction that favours regions in close proximity. The principle of 3C technology is that interactions between distant regulatory regions that come close together in the 3D space will be more frequently detected than random interactions (Dekker *et al*., 2017). Newer technologies coupled to next-generation sequencing (Hi-C, Capture Hi-C) have enabled the detection of topologically associating domains (TADs), defined as local regions within a chromosome with a high density of interactions (contact clusters) that also exhibit insulation from one another (Dixon *et al*., 2012; Nora *et al*., 2012; Sexton and Cavalli, 2015).

TADs are thought to regulate gene expression by increasing the frequency of intra-domain promoter-enhancer interactions and insulating against spurious inter-domain interactions. TADs are formed via cohesin-mediated loop extrusion, whereby DNA is bidirectionally extruded through the ring-shaped cohesin complex until it is halted by convergently oriented CTCF to form a TAD boundary (Golfier *et al*., 2020). It is further assumed that TAD boundaries implicitly prevent transcription read-through and constrain the spread of silencing chromatin. Parameters such as CTCF density and orientation, as well as DNA methylation have been shown to affect TAD direction, size and overall structure. In the last few years, increased sequencing depth and resolution have started to reveal even finer patterns of chromosome contacts and internal sub-TAD structures that can be correlated with functional properties. Examples of finer structures include “dots” associated with loops, and “stripes” that are interpreted as evidence of a dynamic cohesin-mediated “loop extrusion” process whereby some TADs and sub-TADs are formed by the threading of DNA loops through the ring-shaped cohesin complex until it paused by CTCF proteins. HiC techniques have also identified that the higher order structure is further shaped by nucleosome accessibility and divides into A and B compartments, each with distinctive chromatin and transcription features.

Mouse models in which an imprinted locus can be deleted and transmitted through either the male or female germline, have enabled allele-specific Hi-C profiles for the *Igf2-H19*, *Dlk1-Dio3* imprinted loci. For these loci in the mouse, it has been shown that the maternally and paternally imprinted genes are located together in large TADs that are similar in both parental alleles. Within the TADs, differential binding of CTCF creates allele-specific sub-TAD structures that provide the instructive or permissive context for imprinted gene activation during development (Llères *et al*., 2019).

A limitation to studying imprinted genes in humans has been the need for family studies to ascertain the parental origin of genes. Technologies that detect long range cis interactions fortuitously link single nucleotide polymorphism (SNP) variants within a chromosome and provide molecular haplotype information. One of the first studies to use haplotype phasing in Hi-C data from a human lymphoblastoid cell line, GM12878, detected allele-specific long-range interactions between a distal locus, termed HIDAD (Distal Anchor domain) and the promoters of the maternally expressed *H19* and the paternally expressed *IGF2* (Rao *et al*., 2014). GM12878 cells are part of the HapMap project and enabled the assignation of associations to maternal or paternal chromosomes. *IGF2-H19* has been studied in great depth as the archetypal locus for allele-specific interactions (Rao *et al*., 2014; Ye and Ma, 2020). However, the allele-specific-methylation-sensitive-CTCF-binding-for-alternative-looping paradigm as established for *IGF2-H19* is not universally true for all imprinted gene clusters. Imprinted genes occur mostly within clusters, and such domains are interspersed with genes that escape allele-specific silencing.

In this study, we sought to examine how the higher order chromatin conformation structures differ between the active and silent alleles at loci containing genes that are allele-specifically expressed in humans. To this end, we assembled a HiCFlow pipeline for processing raw Hi-C data for haplotype phasing and construction of allele-specific chromatin conformation profiles. A number of existing pipelines, including HiC-Pro (Servant *et al*., 2015), are capable of performing allele-specific HiC. However, these require a pre-phased haplotype as input as they cannot perform *de novo* haplotype assembly from input HiC data. Moreover, most do not have functionality to generate and visualise between-sample normalised differences in contact frequency. As such, we opted to assemble a custom pipeline that integrates the required functionality into a single workflow (see methods, SI Fig 1b). Following this, we were able to characterise allelic differences at human imprinted gene clusters to establish the epigenetic framework for differential association frequencies. We focused on the following three imprinted loci: the *IGF2-KCNQ1* (BWS/SRS locus, chromosome 11p15.5), *SNRPN* (Prader Willi Angelman (PWS-AS) locus, chromosome 15q11-q13), and *DLK1*-*DIO3* (chromosome 14q32.2).

Our analyses indicate that imprinted gene domains are not uniformly organised within a canonical higher order structural profile regulated by elements within the ICRs. At the *IGF2-H19* locus, the ICR plays a direct role in directing allele-specific CTCF-mediated higher order chromatin structures consistent with loop extrusion models, whereas at other loci, the ICR may have indirect or no specific effect. Allele-specific compartmentalisation was observed in some cell lines at the *SNRPN* and *DLK1* loci, and we observed that increased CTCF binding at the *SNRPN* locus occurred with concomitant increased allele-specific contact frequencies.

Rather than remaining spatially and temporally separated from their non-imprinted neighbouring genes, imprinted gene clusters share TADs with non-imprinted genes. Indeed, most allele-specific interactions occur within subTADs. Imprinted domain boundaries may be delimited by TAD structures, but some allele-specific associations can occur across TAD boundaries with concomitant effects of allele-specific expression. Allele-specific interactions were not confined to imprinted domains. In an unbiased genome-wide screen, we detected additional allele-specific TADs (ASTADs). The ASTAD distribution varied between cell lines. We found 8-32% of genes with allele-specific expression to be located within ASTADs. Regions of high genetic variability, such as olfactory receptor loci, the bitter taste receptor (*TAS2R*) gene cluster and the keratin gene (*KRT*) cluster were found to be within ASTADs.

## Results

### A region capture data set with a HiCFlow pipeline provides an effective platform for haplotype phasing of imprinted gene loci

To examine imprinted loci at high resolution we first generated a Region Capture Hi-C (RC-HiC) dataset in a human breast epithelial cell line, 1-7HB2. This diploid cell line has previously been used to examine allele-specific expression and imprinted methylation for several imprinted genes (Nativio *et al*., 2009; Woodfine, Huddleston and Murrell, 2011; Ito, Nativio and Murrell, 2013; Niemczyk *et al*., 2013) and been shown to have methylation and expression profiles consistent with the maintenance of normal imprinting in a somatic cell line. Using a tiled probe RC-HiC approach, combined with a frequent 4 base-cutter restriction enzyme (*MboI*) we generated capture regions (totalling 25Mb) at 6 imprinted chromosomal loci, the largest regions included *SNRPN, DLK1*-*DIO3* and *IGF2-KCNQ1* (capture region IDs: CR_3, CR_2 and CR_4, respectively, SI Table 1). In total 34,399 probes were used covering approximately 4.1Mb (∼16.1%) of the capture regions.

Our RC-HiC dataset yielded approximately 40 million valid read pairs with a mean coverage of almost 1700 read pairs per kilobase and was comparable to published high resolution Hi-C datasets at the same genomic regions (SI Fig 1a). We assembled a Hi-C analysis pipeline (HiCFlow) to process raw Hi-C data to normalised matrices and for haplotype phasing (SI Fig 1b and Methods). This enabled the construction of allele-specific chromatin conformation profiles (alleles designated “A1” and “A2”). Separating the allelic profiles at imprinted loci and visually presenting them as A1 and A2 matrices showed subtle allelic differences in contact frequency. Therefore, we added a subtraction matrix function to the HiCFlow pipeline for highlighting interaction differences between alleles. To further emphasise regions with consistent directional bias, the subtraction matrices were denoised using a median filter (SI Fig 1c). We used the *IGF2-H19* locus to benchmark the allele-specific subtraction parameters and confirmed that the known allele associations could be robustly detected (SI Fig 1c). The subtraction matrix methodology provides a compromise between false positive calls in regions of low interaction density and missing interactions in regions of high interaction frequencies.

The RC-HiC library provided an excellent dataset to test and refine the HiCFlow pipeline. We included regions of non-imprinted genes that could be tested as negative controls. These showed similar profiles when separated into A1 and A2 alleles, and only slight indistinct differences in a subtraction matrix (SI Fig 1d).

### Imprinted gene clusters in human normal breast epithelial cell line exhibit variable patterns of allele-specific associations

We used HiCFlow to define allele-specific association profiles at three imprinted gene clusters in our 1-7HB2 RC-HiC library. These included the *IGF2-KCNQ1* (BWS/SRS locus, chromosome 11p15.5), *SNRPN* (Prader Willi Angelman (PWS-AS) locus, chromosome 15q11-q13), and *DLK1*-*DIO3* (chromosome 14q32.2) loci (Fig 1).

**Figure 1:**
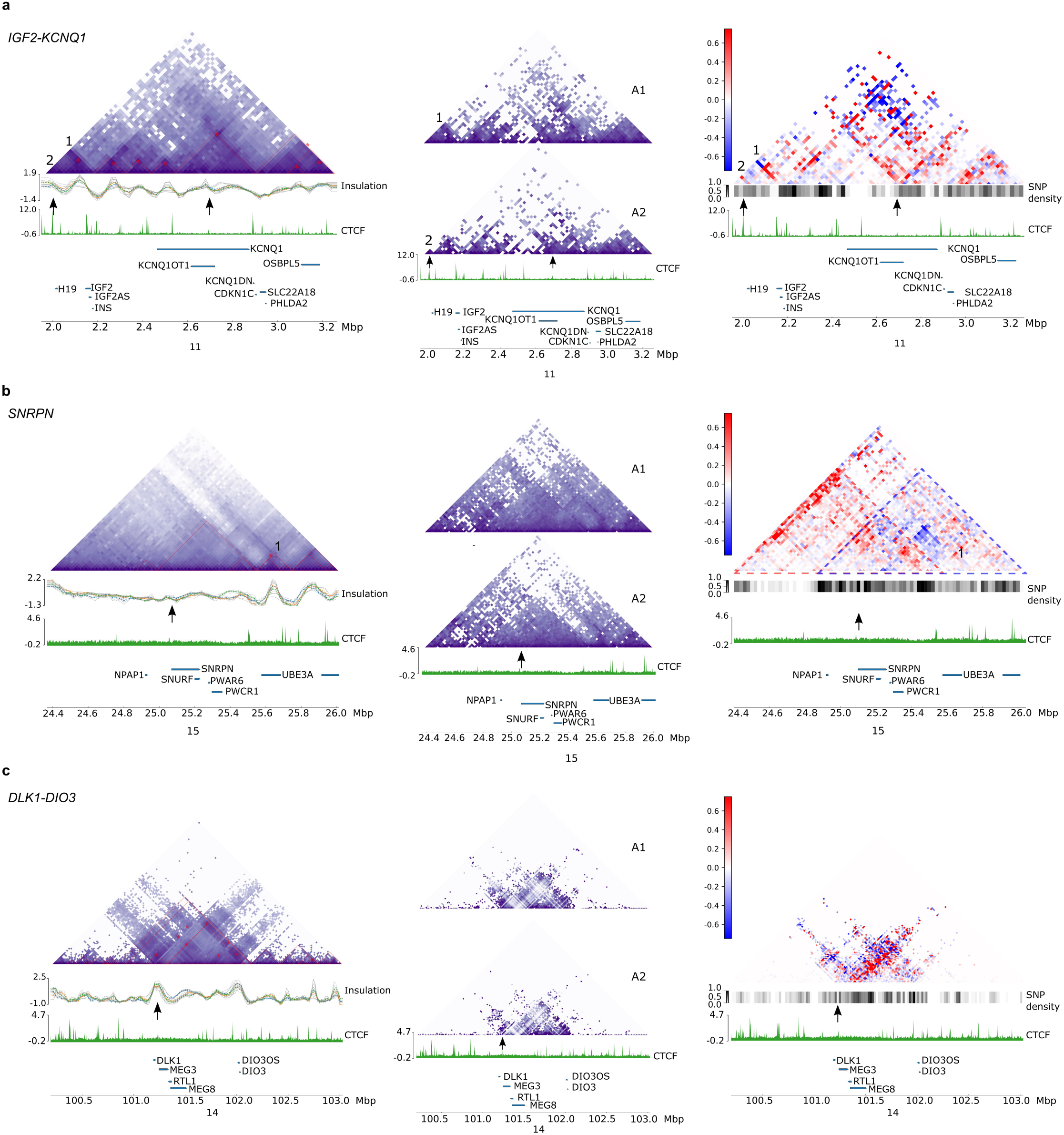
Allele-specific associations at three imprinted gene clusters in a Region Capture HiC (RC-HiC) dataset of human breast epithelial cell line 1-7HB2. **a)** *IGF2-KCNQ1*, with the known *IGF2* and *H19* enhancer interactions labelled 1 and 2. **b)** *SNRPN* locus. **c)** *DLK1* locus. For each locus, the full (diploid) contact matrix (binned at 10kb resolution) is presented, showing the average of all interactions. Below the diploid contact matrix is the TAD insulation score (Ramírez *et al*., 2018), the CTCF track, and the gene track, with imprinted genes annotated. The positions of DMRs with imprinting control functions are highlighted with arrows above the CTCF track for each matrix. Adjacent to the diploid matrices are the haplotype phased allele-specific matrices (A1 above and A2 below) and a subtraction matrix highlighting the differences between the alleles. Enrichment of A1 relative to A2 (blue), enrichment of A2 relative to A1 (red), the scale bar represents distance-normalised differences between A1 and A2. A SNP density track is included to indicate areas of reduced SNP densities that cannot be haplophased. Allelic differences in these regions cannot be called. Coordinates refer to genome build GRCh37/hg19.

Our RC-HiC library captured a 1.2MB section *H19-KCNQ1* domain and included the proximal enhancer to the *H19* gene (Nativio *et al*., 2009), both imprinting control regions (*H19*-DMR and Kv-DMR, marked by arrows above the CTCF track in Fig 1a) and CTCF sites upstream of the *OSBPL5* promoter. It does not include the HIDAD region described by (Rao *et al*., 2014). In Fig 1a, the diploid contact matrix for the region shows a series of dense interactions forming several TADs, against a backdrop of cross-TAD interactions, reminiscent of contiguous TAD cliques (Paulsen *et al*., 2019; Liyakat Ali *et al*., 2021). The *IGF2* gene promoters are located at the TAD boundary on both alleles (Fig 1a). The allele specific matrices display marked differences between A1 and A2, particularly at the *H19*-DMR and the Kv-DMR regions. In the A1-allele, the *H19*-DMR falls within a TAD, whereas in the A2-allele the DMR subdivides the TAD. By contrast, the ICR at the *KCNQ1OT1* promoter region (Kv-DMR), forms a weak subTAD boundary. The difference in allele-specific enhancer interactions with *IGF2* (labelled 1, in Fig 1a) and *H19* (labelled 2 in Fig 1a) is most clearly seen as a blue signal for the A1 allele and a red signal for the A2 allele in the subtraction matrix. This fits with the known shared enhancer model for *IGF2* and *H19* promoters being regulated by the CTCF sites in the *H19*-DMR (Bell and Felsenfeld, 2000; Hark *et al.,* 2000; Szabó *et al.,* 2000; Kurukuti *et al.,* 2006; Engel *et al.,* 2008; Nativio *et al.,* 2009).

The subtraction matrix further indicates a mosaic pattern throughout the region rather than an enrichment of associations of one allele over another such as a complete single coloured TAD. Instead, allele-specific bidirectional stripes from the boundary at the *IGF2* gene and KvDMR regions are evident. At short distances these stripes show a bias towards the A2 allele, whereas at longer range the bias is towards the A1 allele. At the KvDMR the A2-allele has bidirectional interactions towards the CTCF sites upstream of the *KCNQ1* and CTCF site within the *KCNQ1* gene, as well as towards the CTCF sites at the *KCNQ1DN, CDKN1C* and *PHLDA2* genes.

At the *SNRPN* locus, our RC-HiC library captured a 1.6MB region including *NPAP1* to *UBE3A*. The bipartite ICR (marked with an arrow in Fig 1b) is flanked by two CTCF sites. Neither of these sites forms a strong TAD boundary as seen by the weak insulation score below the matrix in Fig 1b. The strongest insulation score in the region is at the *UBE3A* transcription unit corresponding to a small TAD. Overall, TADs are weak/not clearly defined, which could be explained by the low number of CTCF binding peaks at the locus. When the matrix is split into A1 and A2 profiles according to the phased haplotypes, the A2 allele has a fewer long-range associations. The subtraction matrix shows that the region has directional allelic biases: most A2-allele associations (red) are towards the left, whereas A1 (blue) are towards the right. We have highlighted these as triangles with dotted outlines (Fig 1b). The scarcity of CTCF binding, leads us to postulate that the TAD-like structures at this locus, may be formed through phase condensation of heterochromatic compartments. However, region capture data is not suitable for analysis of compartments by current available methodologies.

The 1-7HB2 RC-HiC library captured a region of ∼ 2.5MB around the *DLK1-DIO3* locus. In the diploid contact matrix, we identified a TAD domain overlapping the imprinted region (Fig 1c). The DMRs (both IG-DMR/*MEG3*-DMR, marked by an arrow) are within this TAD and appear to form a subTAD boundary (Fig 1c). However, at this resolution it is also likely that the subTAD boundary is formed by a CTCF site upstream of *DLK1*. In the individual allele matrices, the A1-allele forms a slightly larger subTAD with CTCF bound region upstream of the *DLK1*, while on the A2-allele, the subTAD is more clearly anchored at the ICR (Fig 1c). The subtraction matrix shows a V-shape above the DMR. Towards the *DIO3* locus it forms a predominantly red A2-allele stripe whereas upstream of *DLK1* it forms a blue A1-allele stripe.

It has been proposed that ICRs such as the above DMRs, mediate their epigenetic functions by directing allele-specific chromatin conformation. However, not all imprinted genes contain CTCF sites at their ICRs (Woodfine, Huddleston and Murrell, 2011). Allele-specific chromatin conformation in the 1-7HB2 cell line for the above imprinted gene clusters, indicate that while the *IGF2* locus is very clearly shaped by the *H19*-DMR (ICR) that regulates CTCF site availability, this is not invariably the case at other loci. Indeed, at the *SNRPN* locus TAD-like structures are assembled in the absence of CTCF sites, and independent of the ICR.

### Do imprinting control regions directly participate in allele-specific interactions?

To further examine whether known ICRs are anchor points for allele-specific chromatin interactions we conducted allele-specific viewpoint analyses using haplotype phased Hi-C data from following cell lines GM12878, IMR-90 and H1-hESC, alongside our 1-7HB2 library. We compared these to the relevant subtraction matrices, to which we also added Peakachu loops (Salameh *et al*., 2020). Viewpoint analyses enables focused examination of associations with the ICRs in high resolution data sets, unobscured by the intrinsic density of HiC data. Adding the additional cell line data sets enabled us to investigate how expression, methylation and heterochromatin compartments correlate with allele-specific chromatin conformation. EBV transformed lymphoblastic cell lines such as GM12878 retain DNA methylation profiles consistent with monoallelic expression for several imprinted genes (Baran *et al*., 2015). GM12878 has been haplotype phased previously, as one of the original International HapMap Project cell lines. The remaining cell lines are karyotypically normal diploid and the publicly available HiC data have suitable read depth (SI Fig 1a), to suggest that they could be haplotype phased in our HiCFlow pipeline.

We first examined the *IGF2-H19* and the *KCNQ1* loci. In GM12878 cells, the maternal origin of the *H19* interactions with the downstream HIDAD locus and the reciprocal paternal *IGF2*-HIDAD interactions have previously been demonstrated (Rao *et al*., 2014). We used this information to set the paternal interactions as A1 (blue) at the *H19*-DMR and maternal interactions at the *IGF2* promoter regions as A2 (red) in the subtraction matrices for all three cell lines. This enabled us to also assign the parental origin to the nearby *KCNQ1* locus. Figure 2 demonstrates the effects of the ICRs on higher order structures at the *H19* and *KCNQ1* loci. We note that the *H19*-DMR strongly anchors the maternal allele-specific interactions as can be seen in the subtraction matrices (Fig 2a). This association is independent of the *H19* RNA levels which are high in H1-hESCs, and relatively low in GM12878 and IMR-90 (SI Fig 2a). *IGF2* transcript levels in IMR-90 and H1-hESC are higher than in GM12878 cells (SI Fig 2a). Interestingly, the three cell lines vary for the CTCF interactions with the *H19*-DMR, possibly indicating different tissue specific enhancer associations. Both GM12878 and IMR-90 show this ICR associating with HIDAD region (∼0.3Mbp downstream of the ICR), while in H1-hESCs the ICR interacts with regions further downstream (Fig 2a). At the *H19*-DMR viewpoints (Fig 2a, bottom), IMR-90, and GM12878 show peaks of high frequency A2 (maternal) associations, 50-200kb downstream of this ICR, which correlates with *H19*-enhancer sites. *H19*-DMR also forms weaker biallelic associations at sites up to 150kb upstream. H1-hESCs in contrast, has a stronger biallelic association peak upstream of the ICR and fewer allele-specific enhancer peaks downstream, which may reflect less stable imprinted expression previously reported in human ESCs (Rugg-Gunn, Ferguson-Smith and Pedersen, 2007). Overall, despite the variable expression, the subtraction matrices show similar “stripe” structures in all three of the cell lines, which would be consistent with an allele-specific loop-extrusion between CTCF sites. Thus, the *H19* DMR is an anchor point for a scaffold of stable allele-specific associations at the *IGF2-H19* that ostensibly only depend on whether this ICR is correctly methylated.

**Figure 2:**
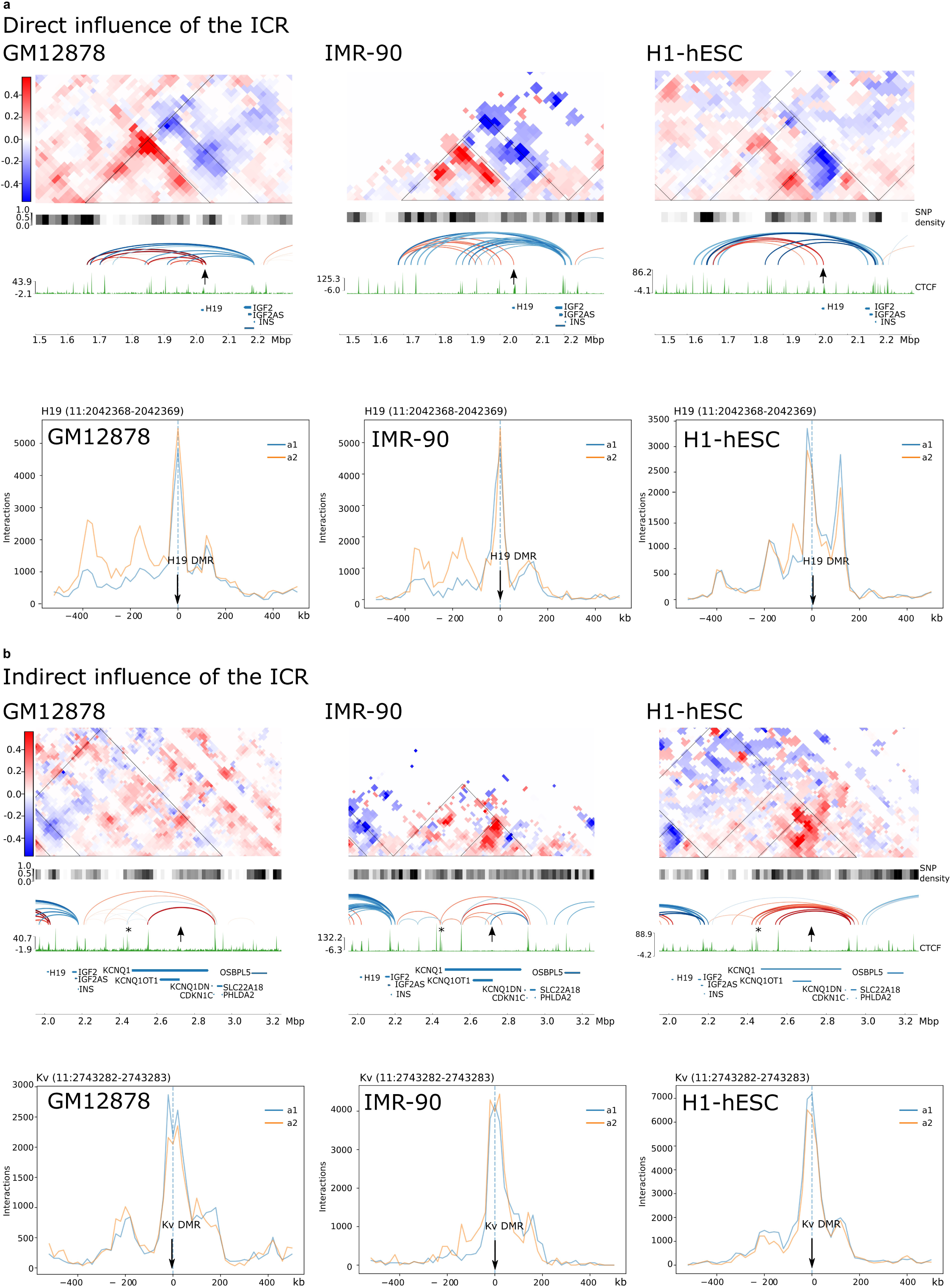
Variable effects of the two imprinting control regions (ICRs) on chromatin conformation at the Beckwith Wiedemann Syndrome locus. **a)** Direct influence of the ICR in structuring local allele-specific chromatin conformation at the *IGF2-H19* locus, in GM12878, IMR-90 and H1-hESC. Denoised subtraction matrices show that the CTCF regulated *H19-*DMR (arrow) subdivides the region (10kb resolution, blue areas correspond to A1, paternal allele, red areas to A2, maternal allele). Below the matrices we have the SNP density, and allele-specific loops, generated by Peakachu (red maternal, blue paternal), that corresponds to maternal and paternal expression of *H19* and *IGF2* respectively. The colour of loops matches the underlying value of the subtraction matrix. The lower panels represent the viewpoint interaction traces for each cell line showing interactions between the *H19*-DMR and loci 400kb in both directions (blue trace A1, paternal, orange A2, maternal). The *H19*-DMR is methylated on the paternal allele in normal cells). **b)** Indirect influence of the ICR in structuring local allele-specific chromatin conformation at the *KCNQ1* locus, in GM12878, IMR-90 and H1-hESC. Subtraction matrices and viewpoint interaction traces as in (a) above but focused on the *KCNQ1* locus and the KvDMR ICR (arrow), normally methylated on the maternal allele. Subtraction matrices, and the allele-specific loops below display variable structural effects on chromatin conformation by the KvDMR. The viewpoint interaction traces mostly show weak biallelic interaction traces for the KvDMR associations. The CTCF site highlighted with an * is “region 3”, previously been shown to be important for allele-specific (maternal) expression of *KCNQ1* (Naveh *et al*., 2021).

In contrast at the *KCNQ1* locus, the ICR (KvDMR) has a weaker and more variable effect on the higher order structure (Fig 2b). Viewpoint analyses at this ICR showed that it formed associations about 200kb upstream and downstream, in all the cell lines tested, albeit weaker than that seen for the *H19*-DMR and not as allele-specific. In IMR-90 cells there is a maternal specific peak (A2-allele) approximately -100kb of the KvDMR (Fig 2b, bottom), that is also seen in 1-7HB2 (SI Fig 2b). For GM12878 and H1-hESC the associations with the Kv-DMR viewpoint are biallelic (Fig 2b, bottom). This ICR is a promoter for *KCNQ1OT1* for which we confirmed the expression in these cell lines as well as in 1-7HB2 (SI Fig 2a). Median levels of methylation at KvDMR are about 50% for IMR-90 and GM12878 (SI Fig 2c) which is consistent with the expected for allele-specific methylation and expression and consistent with our previous reports for 1-7HB2 and IMR-90 (Woodfine, Huddleston and Murrell, 2011). For H1-hESC methylation levels were less than 25% (SI Fig 2c) which is consistent with hypomethylation and biallelic expression of *KCNQ1OT1*. Thus, the viewpoint analysis for the IMR-90 cells fits with allele-specific regulation of loops by the ICR. However, for the other two cell lines there is more ambiguity regarding regulation by ICR.

In all three cell lines the subtraction matrices indicate that associations surrounding *KCNQ1* are predominantly on the maternal allele (Fig 2b). The CTCF signal is weak at the KvDMR in these cell lines (Fig 2b), and there is controversy in the literature about whether this ICR contains CTCF binding sites (Fitzpatrick *et al*., 2007; Lin *et al*., 2011; Prickett *et al*., 2013; Zhang *et al*., 2014; Schultz *et al*., 2015). A paternal-specific loop connecting the KvDMR and other CTCF sites was only present in IMR-90. In IMR-90 the KvDMR is the anchor point of a small subTAD and bidirectional allele-specific loops with CTCF sites in *KCNQ1* (A2, maternal) and *CDKN1C* (A1, paternal) (Fig 2b). We note that in these cell lines, several maternal loops were formed between an intragenic CTCF site within the *KCNQ1* gene (marked with an asterisk in Fig 2b) and other sites at the locus. Recently this CTCF site has been described as “region 3” by Naveh et al (Naveh *et al*., 2021) and it was proposed that interactions between this site and surrounding CTCF sites drives transcription of *KCNQ1* and *CDKN1C* on maternal alleles and is required for normal methylation of the KvDMR. SNPs within this CTCF binding site have previously been reported to be associated with a risk for loss of methylation at this ICR (Demars *et al*., 2014). If this model is correct, then the maternal conformation would prevent *KCNQ1OT1* expression. *KCNQ1OT1* transcription from the paternal allele could potentially displace the intragenic CTCF binding at *KCNQ1* on the paternal allele to reciprocally prevent *KCNQ1* and *CDKN1C* transcription. Thus, unlike the *H19*-DMR, the KvDMR has an indirect effect on chromatin conformation at the locus.

At the *SNRPN* locus, viewpoint analysis at the bipartite imprinting control region, indicate a low frequency of associations within a +400kb window from the ICR (SI Fig 3). GM12878 showed an allele-specific interaction with the PWS AS IC viewpoint at about +400kb, that corresponded with a small A1 enriched TAD-like area on the subtraction matrix. However, allele-specific loops between the ICR and other sites were not identified by the Peakachu algorithm, as shown below the subtraction matrix (SI Fig3, left). We did not detect similar interactions in the viewpoint plots for IMR-90, 1-7HB2 and H1-hESC, despite the subtraction matrices indicating a strong accumulation of allele-specific associations especially in H1-hESC (IMR-90 due to reduced SNP density at this region is uninformative). In the embryonic cells, there is more CTCF binding at this locus, compared to other cells and a more distinct TAD structure. Unexpectedly, despite this strong difference in the subtraction matrices, the methylation data for CpG sites at the ICR indicate that H1-hESC is hypermethylated (SI Fig 2c). *SNRPN* RNA (normally paternally expressed) is present in all these cell lines, but at a lower level in IMR-90. The bipartite imprinting centre at the *SNRPN* locus therefore seems to have an effect on allele specific chromatin conformation at the wider locus. This effect is not reliant on the methylation status of the DMR region within the ICR, at least in H1-hESC cells. The substantial allelic interaction differences in H1-hESC, despite hypermethylation of the DMR may reflect the stability of imprinted gene expression of this locus in embryonic stem cells which occurs independently of DNA methylation (Kim *et al*., 2007; Rugg-Gunn, Ferguson-Smith and Pedersen, 2007).

At the *DLK1-DIO3* locus, only IMR-90 and 1-7HB2 cells showed, allele-specific associations with the IG-DMR viewpoints (SI Fig 4, right). The subtraction matrices for IMR-90, 1-7HB2 and H1-hESC show an allele-specific stripe of interactions from the IG-DMR/*MEG3*-DMR with CTCF sites up to *DIO3* (and beyond in the case of H1-hESC). There may also be a stripe in the opposite direction, however, this is less consistent between cell lines. The *MEG3*-DMR contains CTCF sites, which have been reported to be important in maintaining imprinting in somatic tissues (Kagami *et al.,* 2010a). The strength of allele-specific loops (shown below the subtraction matrices) do not correlate with mRNA levels for *DLK1, MEG3, MEG8 RTL1* or *DIO3* in these cell lines. Indeed, 1-7HB2 and IMR-90 which had the lowest level of expression for these genes, showed the strongest allele-specific loops, and more intense differences between A1 and A2 on the subtraction matrix (SI Fig 4). IMR-90 cells which showed the most distinct difference in allelic conformation was also the only cell line with methylation levels likely to support allele-specific methylation (SI Fig 2c).

The analyses of these four imprinted ICRs indicate that they participate in chromatin conformation to variable degrees in normal cell lines. At the *H19* locus, the *H19*-DMR robustly directs allele-specific chromatin conformation in keeping with a CTCF mediated methylation sensitive enhancer competition model. Here it seems that the chromatin conformation is a stable scaffold even in the absence of *H19* or *IGF2* expression. The IG-DMR seems to direct the chromatin conformation, when normally methylated as seen in IMR-90 cells. At other loci, the ICRs can have indirect effect on chromatin conformation such as the *KCNQ1* and *SNRPN* loci. At the *SNRPN* locus, where there is low amount of CTCF binding, allele-specific associations are present but don’t seem to be driven by the ICR. These results suggest ICRs utilise a variety of mechanisms in addition to CTCF insulation to facilitate allele specific chromatin conformation at imprinted loci.

### Allele-specific compartment differences and effects of imprinting domains on neighbouring loci in normal cells

It is not yet understood how imprinted domains are contained locally and why they do not spread across an entire chromosome. It is expected that chromatin structural elements and compartmentalisation confine imprinted genes to TADs or subTADs to prevent allele specific associations spreading beyond their domains. To examine how far allele-specific associations spread and to detect A/B compartments, we added the CscoreTool (v1.1) (Zheng and Zheng, 2018) to our HiCFlow pipeline and examined a wider 3-6Mb window around each imprinted cluster.

At the *H19*-*IGF2* locus allele-specific interactions did not extend beyond the HIDAD region (chr11:1,500,000) in our cell lines (SI Fig 5a). Interestingly, the recently identified associations between *KRTAP5-6* and *INS* are present within this region (Jian and Felsenfeld, 2021). At the *KCNQ1* locus, we identified looping interactions extending from the *KCNQ1* region to *NUP98* and *RRM1* in an adjacent TAD in H1-hESC. *NUP98* and *RRM1* are both monoallelic in this cell line, but are not known to be imprinted, which suggests that monoallelic interactions can and do extend beyond a TAD containing imprinted genes (Fig 3a, SI Fig5a). The Cscore analysis for this locus indicates that it is located within a 4Mb active A-compartment on both alleles in all three cell lines shown as a red bar below the allele specific matrices in SI Fig 5a.

**Figure 3:**
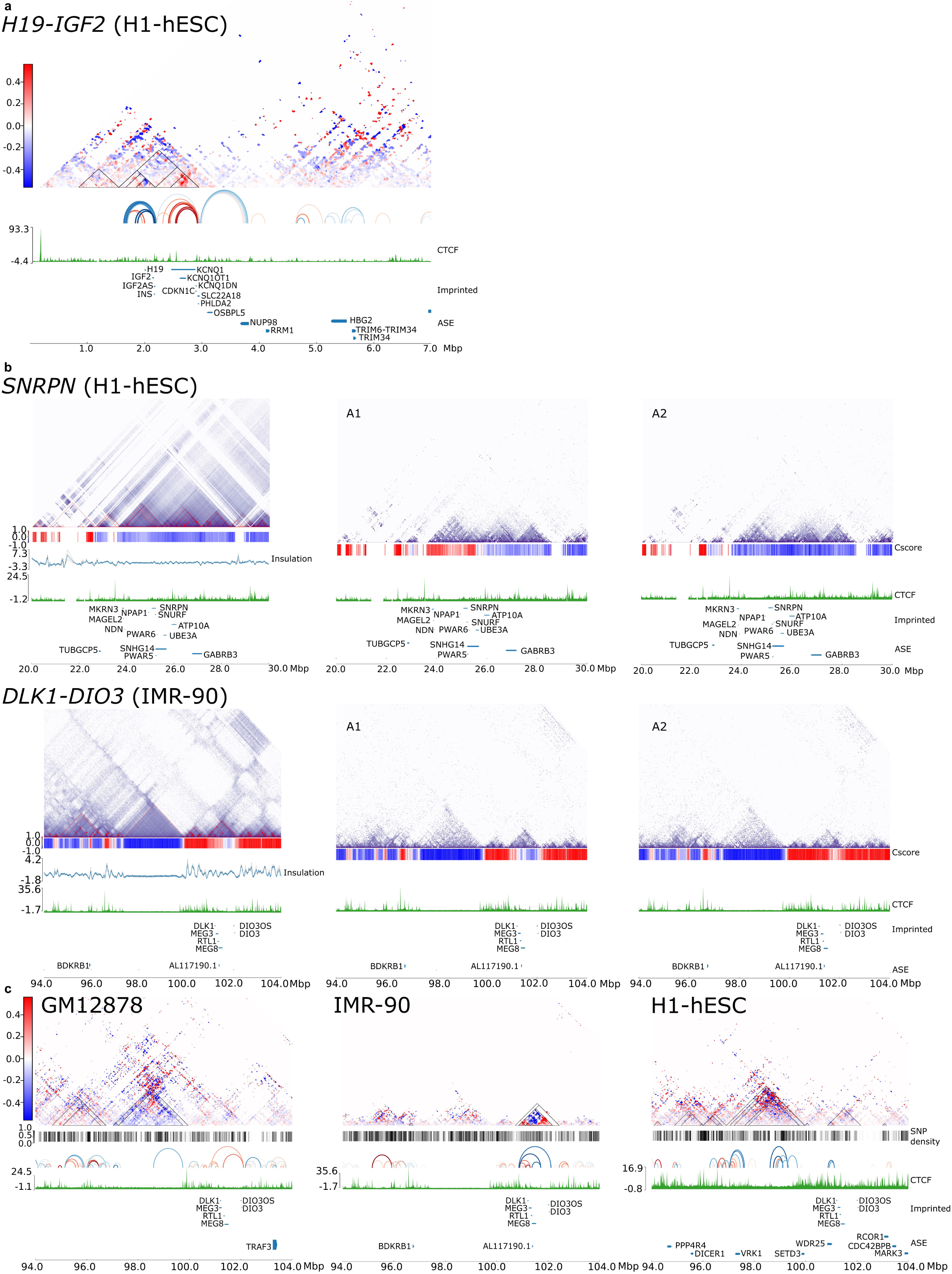
Effects of imprinting domains on neighbouring loci and allele-specific compartment differences in normal cells. **a)** An example of cross-TAD associations from an imprinted gene region. The subtraction matrix at the *H19*-*KCNQ1* locus with allele specific loops in H1-hESC demonstrating cross-TAD association between *KCNQ1* region to *NUP98* and *RRM1* which are allele-specifically expressed (ASE), but not known to be imprinted. Gene density is shown in blue below the CTCF track, with imprinted genes below, and genes with ASE below. **b)** Examples of allele-specific compartmentalisation at *SNRPN* and *DLK1-DIO3* loci. The diploid contact matrix (10kb resolution) with a Cscore below (blue for B-compartment, red A-compartment), followed by TAD insulation score, CTCF track, imprinted genes, and ASE genes. Adjacent to the diploid matrices are the haplotype phased allele-specific matrices (A1 and A2). Note the allele-specific differences in the Cscore track between A1 and A2 alleles at both loci. See SI figure 5 for a comparison of the other cell lines, and subtraction matrices. **c)** Allele-specific cross-TAD associations and additional TAD domains enriched for allele-specific associations near the *DLK1-DIO3* locus. Subtraction matrices, SNP densities, allele-specific loops, imprinted and ASE genes are as described. The *DLK1-DIO3* domain in H1-hESC and GM12878 forms several cross-TAD associations and has weak TAD boundaries. In H1-hESC several genes adjacent to the imprinted domain have allele-specific expression. In GM12878, a nearby TAD (labelled v-TAD) has stronger enrichment for allele-specific associations than *DLK1-DIO3* locus.

The *SNRPN* locus is within a heterochromatin B-compartment, that starts upstream of the imprinted *MKRN3* locus and extends 5Mb towards the telomeric end of chromosome 15 in the three cell lines (SI Fig 5b, and Fig 3b, top). In GM12878 this is a bifold B-compartment that splits into two sections at a point of insulation just after the *UBE3A* gene (SI Fig 5b). Subtle allelic differences are noted in the matrices for the A1 and A2 alleles. In A1, just above the PWS-IC there is a break within the B-compartment which is not present in the A2-allele, which remains within the B-compartment (SI Fig 5b). A similar bifold pattern is seen for this region in IMR-90 cells, except for a region just above the *ATP10A* locus which shows this gene to be in an A-compartment on both alleles in allele-specific matrices (SI Fig 5b). The A2 allele shows further small interruptions in the B-compartment just above the *SNRPN* cluster of genes. The A1 allele does not show these breaks (SI Fig 5b). The most striking difference in allele structure for this locus is seen in the embryonic cells (H1-hESC), where the Cscore analyses returns a similar B-compartment for the full matrix as for the other cells, but in the separate alleles the region above the imprinted genes is distinctly located within a wider active A-compartment in one allele (A1), whereas on the A2-allele the imprinted locus remains in an inactive B-compartment (Fig 3b, top). Interestingly, on the A1 allele, the active compartment seems to spread slightly beyond the boundary upstream of *MKRN3.* H1-hESC seems to have more distinct TAD structures in the separate allele matrices compared to the other cell lines (Fig 3b, top, and SI Fig5b). The subtraction matrices and Peakachu loop algorithm confirm the presence of only a few allele-specific loops in GM12878 and IMR-90 (SI Fig 5b), whereas H1-hESCs have several allele-specific loops, forming a TAD above the *SNRPN* region (Fig 3b). There are also several cross-TAD associations between the *SNRPN* TAD and the adjacent *MKRN3* TAD. These results suggest that the *SNRPN* locus is shaped by phase condensation in conditions of low CTCF binding (Hildebrand and Dekker, 2020), and that when present, CTCF can enhance and stabilise compartmentalisation.

The *DLK1-DIO3* locus, which showed the clearest allele-specific differences surrounding the ICR in IMR-90 (SI Fig 4), was found to have allelic differences in Cscores in these cells (Fig 3b, bottom). The A1-allele of this locus was in a B-compartment whereas the A2 was in an active A-compartment. In GM12878 and H1-hESC cells the locus has no allelic differences in Cscore with both alleles either in a B-compartment (GM12878, SI Fig 6a), or A-compartment (H1-hESC, SI Fig 6a). In IMR-90, the TADs containing the imprinted genes seem to be sharply defined and separate from neighbouring TADs with no overlapping interactions, especially in the B-compartment (Fig 3b, bottom). In H1-hESC, there seem to be more cross-TAD interactions and less sharp TAD borders, although no loops extend from the imprinted region into other TADs. Several allele-specific expressed genes in this cell line are detected in the A-compartment, including *WDR25* and *SETD3* located upstream of *DLK1* in an adjacent TAD (SI Fig 6a, left). This 1Mb sized TAD contains several ncRNAs as well as coding genes, none of which have yet been reported to be imprinted in human, although there is evidence that one or more of the orthologous genes are tissue-specifically imprinted in mice (Tierling *et al*., 2009).

All cell lines have a large 3Mb sized TAD corresponding to a B-compartment that is located 2Mb upstream of the *DLK1* cluster (Fig 3b, bottom, SI Fig 6a). The subtraction matrices show strong allelic bias for predominantly A1 associations in GM12878 cells, and to a lesser extent in H1-hESCs (Fig 3c, right). We have named this the “v-TAD”, after a single coding gene, *VRK1* near the TAD boundary. We examined this region to see whether structural variations were present, specifically duplications that can skew the ratio of allelic associations and found two duplications of 750bp and 89bp (nssv16165643, chr14:98,934,427-98,935,179 and nssv16173610, chr14:97,441,932-97,442,020), that were not associated with any genes and a 304bp duplication in an intron of the *VRK1* gene (nssv16177248, chr14:97,284,401-97,284,704), listed in NCBI data base. In our cell lines, we find no variation in copy number within the v-TAD domain. However, both GM12878 and H1hESC, possess different Indel mutations within the region corresponding to the nssv16165643 duplication. It is unclear to what extent these variants are responsible for allele-specific associations in the v-TAD. Since they do not overlap CTCF sites, we do not anticipate they are responsible for such large-scale allelic changes in GM12878 and H1hESC.

*VRK1* encodes a Serine/Threonine Kinase and is associated with Pontocerebellar Hypoplasia, Type 1A and Microcephaly-Complex Motor and Sensory Axonal Neuropathy Syndromes. It is widely expressed in several tissues and has roles in cell cycle, mitosis and DNA damage responses. It has never been reported to have monoallelic expression. We found it to be highly expressed in all cell lines (SI Fig 6b) and monoallelic in H1-hESC (Fig 3c). The v-TAD region in H1-hESC seemed to form several cross-TAD interactions with adjacent TADs, and further genes (*PPP4R4* and *DICER1*) were found to have allele-specific expression (Fig 3c).

In summary these results indicate that imprinted regions can have allele-specific associations confined within TADs, without differences in compartmentalisation such as at the *IGF2-H19* and *KCNQ1* loci. We have also seen that compartmentalisation can be detected allele-specifically and that imprinted regions when present in active A-compartments, can form looping associations that extend beyond their own TAD regions, with the potential of allele-specifically activating genes outside the imprinted locus.

### Clusters of allele-specific interactions occur throughout the genome as allele-specific TADs

The detection of the above v-TAD prompted us to examine the frequency in which differences in allele-specific associations can be found within TADs genome wide. We therefore performed an unbiased ranking of all TADs to assess the allelic associations differences genome-wide in GM12878, IMR-90 and H1-hESC cells (Fig 4a). We defined allele-specific TADs, (ASTADs) as having higher than expected absolute differences in A1 and A2 associations. TADs containing imprinted genes (SI File 1) had Z-scores of 2.9-5.8 (*IGF2-H19*, in all three cell lines), 5.5-9.1 (*DLK1-DIO3* in IMR-90), and the *SNRPN* locus (2.2-3.1 in H1-hESCs). The v-TAD described above had a Z-score >4 in GM12878 and H1-hESC.

**Figure 4:**
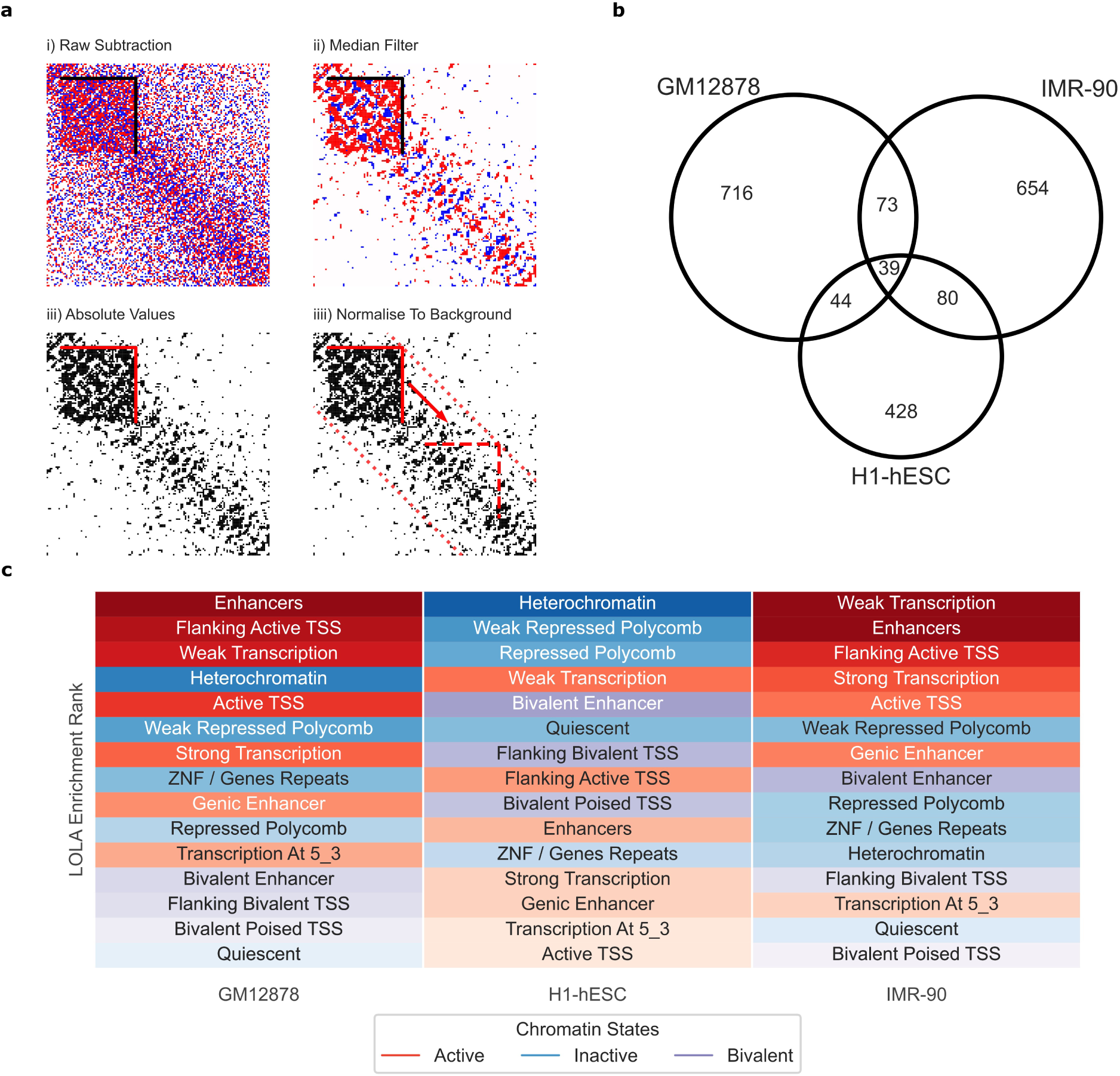
Several TADs genome-wide are enriched for allele-specific associations. **a)** Illustration of ASTAD detection methodology. i) Reference TAD domains are aligned with the raw HiC subtraction matrix. ii) A median filter is applied to remove background noise and emphasise regions of consistent directional bias. iii) The absolute sum of intra-domain allelic differences is calculated. iv) A Z-score is calculated by comparing against the chromosome-wide background level of absolute differences for a domain of equivalent size. TADs with Z-score > 2 are considered ASTADs. **b)** Venn diagram of conserved ASTADs between cell lines. Conserved ASTADS were defined as any set of domain intervals, between cell lines, that shared 90% reciprocal overlap. **c)** ASTAD enrichment across different chromatin states. Chromatin states are ordered, per cell line, according to their enrichment level as determined by LOLA (max rank). ASTADs possess contrasting enrichment characteristics in H1-hESCs compared the differentiated cell lines. IMR-90 and GM12878 were significantly enriched in active chromatin relative to all TAD domains. H1-hESC were enriched for inactive and bivalent states.

Each cell line had its own unique profile of ASTADs distributed across the genome (SI Fig7a and b), although there was a small overlap between cell lines (Fig 4b). Allelic imbalances due to abnormalities resulting in trisomy show up as large blocks of allelic bias, and this was found at the 11q chromosomal region in GM12878 cells (SI Fig7c). Imbalances due to monosomy will be masked as these regions can’t be phased. Random monoallelic effects such as X-inactivation are not expected to show up as ASTADs because most normal tissues have a 50% mix of cells with either of the parental X-chromosomes inactivated. Indeed, in IMR-90 cells where X-inactivation is random, no allelic bias was found on the X-chromosome (SI Fig7d). GM12878 cells, in comparison has skewed X-inactivation (Rao *et al*., 2014), and a large number of allele-specific associations on the X chromosome (SI Fig 7d). The H1-hESC cell line is male and thus these cells do not register heterozygosity for phasing on the X chromosome. A full description of all ASTADs detected is available in SI File 2. Overall, ASTADs vary in size comparable to non-ASTADs (SI Fig 7e).

To determine whether ASTADs were associated with specific chromatin (heterochromatin compartments, DNAse I sensitivity, histone signatures associated with regulatory elements) or genomic (heterozygous SNPs, lncRNA, CpG islands) features, we carried out locus overlap analysis (LOLA) (Sheffield and Bock, 2016) for 15 states previously imputed using chromHMM (Ernst and Kellis, 2012). This revealed that the ASTADs had different characteristics in H1-hESCs compared the differentiated cell lines. IMR-90 and GM12878 are significantly enriched in active chromatin relative to all TAD domains (Fig 4c). H1-hESC were enriched for inactive and bivalent states.

CNVs are known to influence chromatin architecture (di Gregorio *et al*., 2017) and therefore the detection of ASTADs. CNVs may also influence mappability to the reference genome and introduce artefacts in variant discovery and haplotype phasing. Although HiCFlow implicitly removes CNV driven biases through HiCcompare, we carried out an independent analysis of CNV using QDNAseq (Scheinin *et al*., 2014) to assess whether ASTADs were associated with non-normal copy numbers. With the exception of a low magnitude, but significant, enrichment of gain CNV in IMR-90, non-normal CNV regions were not substantially over-represented in ASTADs relative to TADs (SI Table 5 and 7).

### Conserved ASTADs

The three cell lines tested came from three different individuals and represent three different cellular lineages (lymphoblastic, foetal lung and embryonic stem cells). Based on these genetic and the tissue differences, we expect only a few ASTADs would be common to all three cell lines. From the above analysis, we found 39 conserved ASTAD. We hypothesised that common ASTADs would fall into two categories. The first category being sequence directed, such that ASTADs present at the same location in the three cell lines, would share identical SNPs variants. The second category being ASTADs with stable epigenetic mechanisms directing allele-specific chromatin conformation as in genomic imprinting.

We first tested whether conserved ASTADs boundaries possessed the same, i.e. identical genetic variants, across all three cell lines that may influence chromatin structure in a consistent manner. Randomisation testing with repeat random sampling (n = 10,000) of conserved TAD versus ASTAD boundaries (see methods, SI Table 8), indicated significant enrichment (Z-score = 2.13, p = 0.016) of identical genetic variants at conserved ASTAD boundaries compared to conserved TADs.

A similar analysis, examining the distribution of allele-specific methylation (ASM) within conserved TAD versus ASTAD boundaries was performed for each cell line. We did not detect any significant enrichment of in H1-hESC and IMR-90. Enrichment was marginally higher in GM12878 (Z-score = 1,67, p = 0.048). These results suggest that conserved ASTADs are primarily sequence directed due to sharing similar haplotypes.

The conserved ASTAD with the highest ranking allelic difference (Z-score = 4.9 – 9.7), was identified at chr3:195,270,000 – 195,730,000 (SI File 3). However, this region was found to overlap an ENCODE Blacklist region usually associated with anomalously high read mapping signal (Amemiya, Kundaje and Boyle, 2019), possibly due to unannotated repeats in the reference gene sequence. We therefore cannot exclude this ASTAD being an artefact. The next highest ranking conserved ASTADs (chr12:52,530,000-53,390,000, Z-score = 3.6 – 5.3 and chr7:2,910,000-4,810,000, Z-score = 3.6 – 5.6), are not within blacklisted regions.

The ASTAD at the chr12:52,530,000-53,390,000 region contains a cluster of keratin type II cytoskeletal orthologues, involved in hair and epithelial keratin synthesis, and a high association with disease associated variants. This region has been described as an EAFD locus (genetic variants with extreme allele frequency differences) and a feature of such loci are that the SNPs have longer linkage disequilibrium (LD) ranges than random SNPs (Sulovari *et al*., 2017). *KRT1* has been reported to be expressed allele-specifically as a result of cis-regulatory polymorphisms (Pant *et al*., 2006), and a more in-depth analysis has shown allele-specific expression is a complex trait of multiple SNPs having a cumulative effect on gene expression (Tao, Cox and Frazer, 2006). In GM12878, IMR90 and H1-hESCs, none of the *KRT* genes were listed as having allele-specific expression, despite this region showing strong allelic differences in association frequencies (SI Fig 8).

### Imprinted genes in ASTADs

Only 5 Imprinted genes are present in conserved ASTADs. Four of these (*H19*, *IGF2*, *IGF2-AS* and *INS*) are part of the same imprinted gene cluster on chromosome 11, the other is the recently identified pseudo-gene *ATP5F1EP2* (Santoni *et al*., 2017) on chromosome 13 (SI File 1). The *H19-IGF2* cluster is the most studied of imprinted genes, and perhaps this is due to its robust and stable CTCF-mediated imprinting mechanism. Little is known about imprinting mechanisms for *ATP5F1EP2.* However, a close look at this locus showed allele-specific expression of further genes within this ASTAD, including *POLR1D*, *MTIF3*, *USP12* in H1-hESC, and *RPL21* in GM12878. These genes have not been reported to be imprinted. Gene mutations in *POLR1D* underlies autosomal dominant inheritance of Treacher Collins Syndrome (TCS, OMIM 154500).

In our targeted analyses we found that the *KCNQ1, SNRPN and DLK1* loci had several differences between the cell lines and therefore unlikely to be within conserved ASTADs (SI File 1). We screened 115 genes reported to be imprinted in humans (Ginjala, 2001), to determine if they were within ASTADs in any of the cell lines as opposed to being within a conserved ASTAD (SI File 1). In H1-hESC, 45 (39%) of the imprinted genes are in ASTADs. IMR-90 and GM12878 have 38 (33%) and 42 (37%) respectively.

Randomisation testing (see methods) revealed that Imprinted genes were significantly enriched (p <= 0.001) within ASTADs compared to non-ASTADs (SI Table 9). However, the observation that so few imprinted genes were in conserved ASTADs suggest that somatic differences in imprinted gene expression, methylation, and other epigenetic effects, influence the density of allele-specific interactions such that their TADs do not consistently meet the threshold of an ASTAD.

### Allele-specific gene expression in ASTADs

ASTADs are domains with high frequency of allele-specific contacts. Therefore, we examined whether genes located within ASTADs have allele-specific expression and downloaded RNAseq ASE data for GM12878 (3099 bi-allelic, 480 mono-allelic) (Workman *et al*., 2019) and IMR-90 (409 mono-allelic) / H1-hESC (2398 mono-allelic). Only the GM12878 dataset included genes with confirmed biallelic expression. In GM12878, 153 of 480 (32%) of ASE genes were found to be within ASTADs. For IMR-90, this was 73 out of 409 (18%) and for H1-hESC, it was 182 out of 2398 (8%). Randomisation testing (see methods) revealed that ASE genes were significantly enriched (p < 0.001) within ASTADs in GM12878 and IMR-90, but not in H1-hESC (GM12878 (Z-score = 4.64, p = 1.7e-6), IMR-90 (Z-score = 3.43, p = 0.0003), H1-hESC (Z-score = -1.78, p = 0.962).

It has been suggested that polymorphisms within enhancers are more likely to disrupt chromatin architecture and influence gene expression. We therefore assessed whether the heterozygotic variation in enhancers (using public available data from (Nasser *et al*., 2021)) are enriched in ASTADs and found a significantly higher than expected proportion of heterozygous variants in enhancers overlapping ASTADs (chi-square test, p < 0.001) in GM12878 and IMR-90, but not in H1-hESC. In addition, we find that enhancers associated with ASE genes are significantly over-represented in ASTADs (chi-square test, p < 0.001) in GM12878 (SI File 4).

We further found that a cluster of *TRIM* genes on chromosome 11, contained allele-specific (paternal allele) expressed genes (*TRIM5*, *TRIM22* in GM12878, *TRIM6, TRIM6-TRIM34* in IMR-90 and *TRIM34*, *TRIM6-TRIM34* in H1-hESC) and were within an ASTAD in GM12878, and in IMR-90 (SI Fig 5). The ASTAD containing the *TRIM* cluster also contains a cluster of olfactory receptor genes (*OR52 -OR56*), which typically express only one allele, but did not feature in the lists of ASE in these cell lines. Olfactory genes have been shown to form interchromosomal associations and aggregate in foci within the nucleus when they are repressed, with the expressed allele localised outside of such foci (Clowney *et al*., 2012; Markenscoff-Papadimitriou *et al*., 2014).

One ASTAD region of interest included the *TAS2R* gene cluster that encodes an array of Bitter Taste Receptor genes on chromosome 12p13.2. The *TAS2R* gene cluster overlaps an ASTAD in both GM12878 (chr12:10,915,000 - 11,405,000) and in IMR-90 (chr12:10,900,000 - 11,370,000) corresponding to an overlap of approximately 90%, but this may be due to a lack of SNP density in IMR-90 (Fig 5). The ∼400kb ASTAD seems to originate from a weak CTCF binding site as a subTAD within a 800kb wide CTCF defined TAD. We further found that the region had allele-specific differences in Cscores, such that in H1-hESC, one allele was more enriched for the A-compartment, while the other was divided into several smaller A and B-compartments (Fig5a, right). In GM12878 there was allelic variation within A compartments, whereas in IMR-90 there was allelic variation within B-compartment. *TAS2R* genes have not previously been identified as imprinted or to have allelic expression. We examined RNA transcript levels for several of the *TAS2R*s genes at the locus by PCR and confirmed that these were expressed in all three cell lines (Fig 5b). These genes are usually very small single exon genes and despite being in a region of high sequence variability, most of the SNPs are intergenic. Even the lncRNAs, of which there are several at the locus have very small exons and thus we found no informative SNPs to enable us to verify whether genes at this locus have monoallelic expression. Taste receptors, like olfactory receptors are G-protein coupled receptors and they may similarly have monoallelic expression due to allelic exclusion. We examined the *TASR2* clusters on chromosomes 5 (*TAS2R1*) and 7 (*TAS2R3-TAS2R38)* and found that these did not overlap ASTADs. However, to our knowledge this is the first time that taste receptors on chromosome 12 have been reported to be present in an allele-specific chromatin conformation.

**Figure 5:**
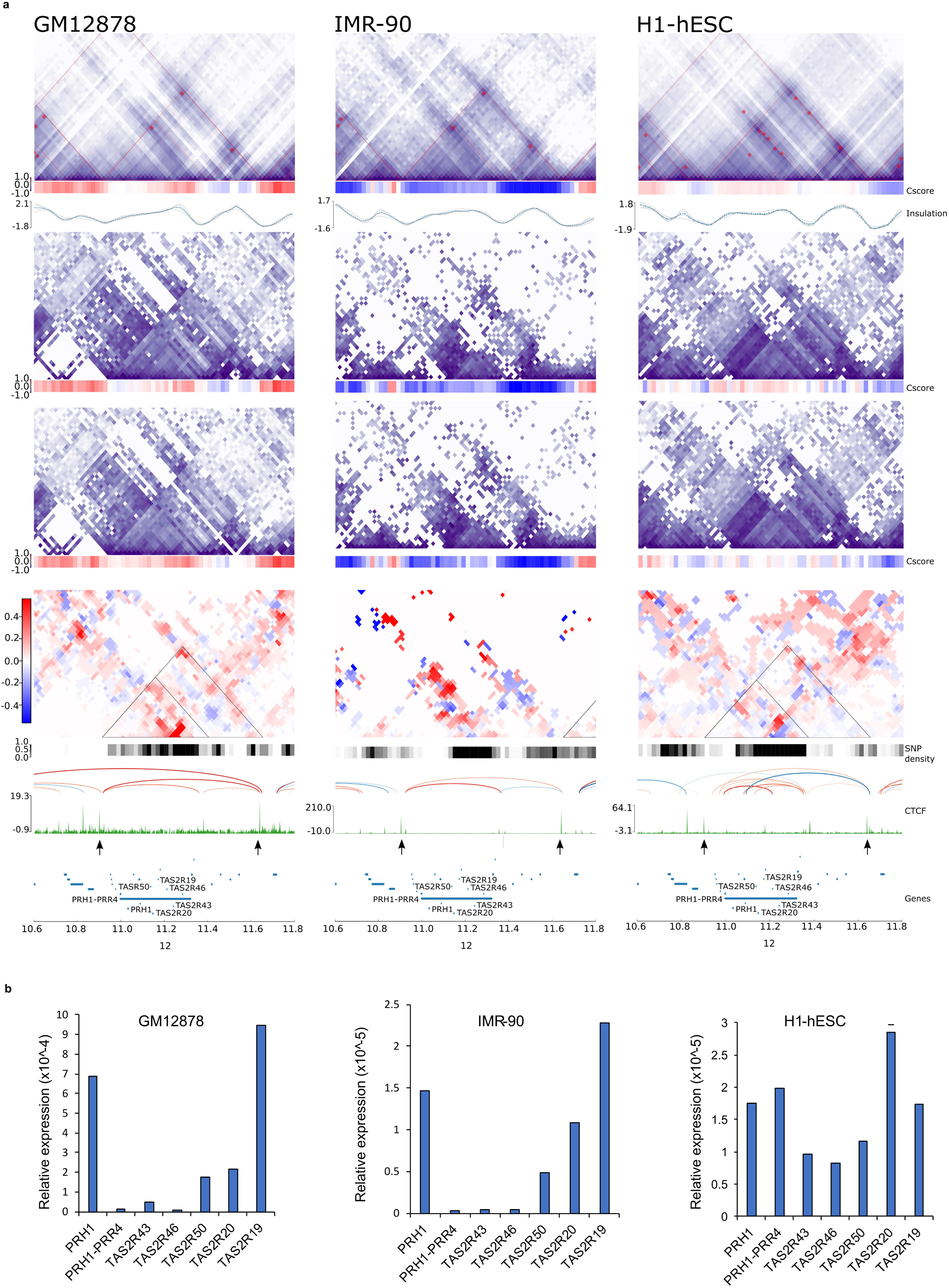
The Bitter Taste Receptor (TAS2R) cluster on Chromosome 12p13.2 is within an ASTAD. **a)** Contact matrices for diploid, haplotype phased alleles and their subtraction matrices at 10kb resolution in GM12878, IMR-90 and H1-hESC. Arrows below the CTCF track indicate boundaries of the ∼800kb TAD which hosts the ASTAD (as a subTAD). ASTAD shown as triangles. The ASTAD is identified in GM12878 (chr12:10,915,000 - 11,405,000, Z-score = 4.06) and IMR-90 (chr12:10,900,000 – 11,370,000, Z-score = 4.20) with >90% overlap. H1-hESC was not called as an ASTAD (Z-score = 1.56). Cscores, below the contact matrices show clear allelic differences, that varied between cell lines. **b)** Quantitative RT-PCR analysis of RNA transcript levels for TAS2Rs genes and the lncRNA *PRH1-PRR4*.

## Discussion

Allele-specific gene expression is a common event in normal cells but can be cause of congenital and acquired diseases including cancer (PCAWG Transcriptome Core Group *et al*., 2020; Przytycki and Singh, 2020). True monoallelic gene expression where one allele is preferentially silenced such as in genomic imprinting, relies on epigenetic mechanisms that initiate and maintain the silencing. In most cases, allele-specific expression is driven by sequence variants located within gene regulatory elements to confer allele-specific preference for transcription factor binding. Genome wide association studies (GWAS) have linked variants and diseases and have enabled insights into complex-trait genetics and important biological processes in gene regulation and mechanisms underlying disease. Typically, these variants (93%) occur in the non-coding intergenic regions, introns, and long terminal repeats (Maurano *et al*., 2012). The major challenge of GWAS lies in interpreting the involvement of these non-coding variants in the aetiology of diseases. Methods that integrate GWAS data with expression quantitative trait loci (eQTL) data to identify associated genes (Zhu *et al*., 2016), and approaches that combine epigenetic data such as DNA methylation (Giambartolomei *et al*., 2018), ChIP-seq, DNase I hypersensitivity have been used to suggest functional hypotheses for variants associated with diseases (Bryois *et al*., 2018). More recently, chromatin interaction information has been used to link GWAS variants to target genes (Javierre *et al*., 2016; Gorkin *et al*., 2019; Yu, Hu and Li, 2019) and more tools are being developed to predict the functional effects of variants in disease including combining artificial intelligence and deep learning with HiC data (Meng, Xiao and Deng, 2021). A systematic study of the genome wide distribution of disease-associated SNPs indicate that they are statistically more likely to be located within TAD borders where they can alter TAD structures (Jablonski *et al*., 2021). Aside from allele-specific differences at imprinted loci, the occurrence of allele-specific TAD structures and their association with allele-specific gene expression has not been extensively documented at a genome-wide level.

In this study we assembled a novel bioinformatics pipeline, HiCFlow, which combines variant calling and haplotype phasing with allele-specific HiC analysis to enable robust investigation and visualisation of allele-specific associations. We initially focused on three imprinted gene clusters that exemplified imprinted domains known to be allele-specifically regulated. The canonical boundary model exemplified by *IGF2*-*H19* is consistent with an allele-specific loop extrusion model, which shows up as a pair of parallel stripes in a subtraction matrix. In this case, CTCF sites anchored at the HIDAD locus (or other sites downstream in a tissue specific manner) interact with CTCF sites at the *H19*-DMR on the maternal allele, and with CTCF sites near the *IGF2* promoter on the paternal allele, when the *H19*-DMR is methylated. The discovery of HIDAD locus as the furthest CTCF anchor point at the boundary of the TAD that contains the whole imprinted *H19-OSBPL5* domain indicates that *IGF2* or *H19* can gain access to a series of enhancers located near to 9 different intervening CTCF pause-sites during loop extrusion. Allele-specific interactions between the HIDAD and *H19* were not detected in H1-hESCs, which may explain previous reports of unstable imprinting of *IGF2-H19* in these stem cells (Rugg-Gunn, Ferguson-Smith and Pedersen, 2007). However, the differential stripe pattern at the *H19*-DMR in all cell lines tested, indicates that this ICR determines the allele-specific chromatin scaffold. In contrast, the adjacent imprinted gene cluster, clearly shows that the ICR (Kv-DMR) regulating imprinting at the *KCNQ1* locus, has weak if any effects on 3D chromatin structure. If there is allele-specific loop extrusion, this occurs at a CTCF site previously identified as “region 3” and shown to be required for setting up a maternal allele-specific loop to facilitate *KCNQ1* expression (Naveh *et al*., 2021). The paternally expressed *KCNQ1OT1* lncRNA transcript is regulated by the Kv-DMR, and its transcription may disrupt CTCF binding at region 3 on the paternal allele to prevent this loop. RNA polymerase 2 has been reported to displace CTCF occupancy before (Lefevre *et al*., 2008), we have evidence that this may be the case at the *KCNQ1* locus, and even if it did, the ICR would have an indirect effect on 3D structure at this locus.

Phase condensation models have not previously been tested at imprinted gene loci. A large body of data from chromatin immunoprecipitation for post-translational histone modification profiles at imprinted loci in mouse and humans (Farhadova, Gomez-Velazquez and Feil, 2019) predict that the silent allele should be in a heterochromatic configuration. Early studies using fluorescent in-situ hybridisation at the *SNRPN* locus demonstrated that active and silent alleles occupy different nuclear compartments (Mahy *et al*., 2002). Adding the Cscore to the HiCFlow pipeline enabled us to confirm allele-specific differences in compartments at imprinted loci. These differences were undetectable at the *H19-IGF2* and the *KCNQ1* loci in these cell lines. Strong differences in allele-specific associations were evident in the subtraction matrices, where strong differences in compartmentalisation was observed, for example at the *DLK1* locus in IMR-90 cells. This correlated with ICR methylation levels that would be consistent with normal imprinting. A striking difference in compartment structure was detected at the *SNRPN* locus in H1-hESC. This was also apparent at this locus in the other cell lines, albeit not as markedly or as widespread as in the stem cells, where the imprinted expression is more stable. These cells had clear evidence of CTCF occupancy at the locus. Current models suggest that CTCF stabilises TAD structures and that compartmentalisation processes counteract the formation of TADs. Our observations at the *SNRPN* locus suggest that CTCF stabilised higher order structures that may be initiated by phase condensation-mediated compartmentalisation.

Within imprinted domains, there are also non-imprinted genes that escape the effects of neighbouring gene silencing and we do not yet understand how imprinting is contained locally and why it doesn’t spread across an entire chromosome. By examining the loci from a wider perspective, we found that each locus occurred within a larger TAD structure, which is similar to observations made by the Feil group in mice for the *Igf2* and *Meg2* loci (Llères *et al*., 2019). The loci we examined suggest that imprinted genes are present within interacting subTADs with weak boundaries. We showed evidence that allele-specific looping associations in imprinting domains that are present in A-compartments can extend to neighbouring TADs. Such observations are rare, and it is feasible that this “neighbour effect” requires that the non-imprinted neighbour is already in an active state, and able to be allele-specifically upregulated by associations with enhancers within the imprinted domain.

Our novel HiCFlow pipeline is suitable for processing multi-sample Hi-C, RC-HiC and Micro-C datasets. It is implemented as a user-friendly Snakemake pipeline and, to our knowledge, is the first workflow to combine haplotype phasing with Allele Specific-HiC analysis. Testing it on a RC-HiC dataset provides proof-of-concept data for a diagnostic assay that can be used to detect allele-specific chromatin conformation in humans with imprinting disease. In this capture data set we only examined a small set of imprinted gene clusters, but these could be expanded to the full set of imprinted genes and would still provide the required read depth when sequenced in-house on standard sequencing platforms. A limitation of our RC-HiC analysis was that we focused on the core regions of the imprinted domains, which enriched for the detection of shorter-range interactions. We would recommend utilising tools, such as CHiCANE and Peaky, to detect long-range interactions (Eijsbouts *et al*., 2019; Holgersen *et al*., 2021). For the purposes of this study, our focused analysis was sufficient to show that different chromatin conformation structures are present at imprinted loci, rather than a “standard ICR centred” structure. In the last decade it has been shown that a subset of patients diagnosed with an imprinting disorder, have multi-locus imprinting disturbances (MLID), characterised by loss of methylation at multiple imprinted loci across the genome (reviewed by Monk *et al.,* 2019). RC-HiC and HiCFlow will also be useful tools in the comprehensive and integrative analysis of MLID.

To investigate the properties of genome-wide allele-specific interactions, we scored each TAD based on the absolute allelic differences observed. The set of top-ranked TADs, denoted “allele-specific TADs (ASTADs)”, were assessed for over-representation of various genomic annotations relative to non-ASTADs. Most strikingly, ASTADs were found to be enriched with polymorphic variants. This is unsurprising since heterozygous SNPs are necessary to distinguish alleles during allelic assignment of read pairs. However, enrichment of INDELs in ASTADs, which were not used for allelic assignment, suggests that high genetic variability plays a role in influencing allele-specific chromatin conformation. Indeed, regions of high variability are prone to allele-specific gene expression as demonstrated by the large body of GWAS studies, that correlate genetic variants with allele-specific binding of transcription factors, DNA methylation patterns and gene expression (reviewed (Wang, Lou and Wang, 2019)). This is further supported by our finding that heterozygous variants are more likely to be associated with an ASTADs if they overlap a known enhancer.

We also found that allele-specific expressed genes were significantly over-represented in ASTADs in GM12878 (32%) and IMR-90 (18%), although not in H1-hESC (8%). Despite enrichment, only a small absolute proportion of ASE genes overlapped ASTADs. In some cases, it is likely that short-range allele-specific interactions may be indistinguishable at the 20kb resolution of our data. In addition, the absence of informative SNPs can prevent ASE detection and detection of allele-specific chromatin interactions.

As expected, imprinted genes were significantly over-represented in ASTADs compared to non-ASTADs. However, imprinted genes did not invariably overlap ASTADs and different chromatin conformation strategies were employed at different imprinted domains. Only the *H19/IGF2* locus was consistently identified as an ASTAD and displayed the same differential looping interaction at the *H19*-DMR across all cell lines. Our cell lines also varied in their stability for imprinted expression at various loci. Allele-specific associations at imprinted loci, also did not correlate with levels of detectable transcripts. It has been suggested that the chromatin conformation forms a scaffold upon which transcription factors can dock and activate gene expression (Ray *et al.,* 2019). Such a scaffold would therefore not correlate with expression levels if the right transcription factors aren’t present. However, while this may be true for some loci, given the overall conserved CTCF binding across multiple tissues and cell types, there is enough variation in TAD structures, and reported experimental evidence indicating that active transcription modulates higher order chromatin structures (Li and Reinberg, 2011). A lack of correlation between looping associations and transcripts, could therefore be due to the post transcriptional effects that affect RNA stability. The strength of chromatin loops between an enhancer and promoter and transcription factor binding kinetics has recently been correlated with transcriptional bursting (Yokoshi, Segawa and Fukaya, 2020; Ma *et al*., 2021). Thus, the frequency and length of time that a gene promoter and enhancer interact affects the frequency and length of a transcriptional burst. Technologies in which such models can be tested experimentally on a genome-wide scale are not yet available.

We were able to confirm that regions of high variability and known to have allelic imbalances in expression such as the olfactory receptor genes were within ASTADs. We were intrigued to find that the bitter taste receptor (*TAS2R*) clusters were present within ASTADs. To our knowledge these genes have not been reported to be subject to allelic exclusion. They are however present in regions of high genetic variability and similar to olfactory receptors. The *TAS2R* family of receptors similar to olfactory receptors are G protein coupled receptors. There are about 25 functional *TAS2R* genes and 11 pseudogenes spread between chromosomes 5, 7 and 11. Extensive population studies have utilised the high variation to examine evolutionary origins of different haplotypes and to identify the selection pressures that an ability to distinguish bitter toxic substances has had on the genetic evolution of this gene family. The functional effects of the variants on G-protein receptor protein structures and the mechanisms whereby they convey taste perception to the brain has been elucidated. However, limited information on their transcriptional regulation exists. It has been recently shown that *TAS2R* genes are not only expressed on the tongue but that they are more widespread and present in heart and respiratory epithelia as well as in the gut and that they may have further sensing functions unrelated to bitter taste. An in-situ hybridisation study has indicated that humans co-express a heterogeneous mix of between 4-11 taste receptors per cell in papillae of the tongue. Although, we do not know if this due to allele-specific expression, our data indicate that this may be the case as ASTADs are commonly associated with allele specific expression.

Overall, this study highlights how genetic sequence variation and the regulatory mechanisms behind allele-specific gene regulation culminate in widespread allelic differences in chromatin organisation that are not confined to imprinted gene loci.

## Methods

### Cell lines

Human mammary epithelial cells (1-7HB2 cell line) were cultured in RPMI-1640 (Sigma-Aldrich), supplemented with 5% fetal bovine serum (Sigma-Aldrich, B6917), 10 ml/l penicillin-streptomycin solution (Gibco, 15140122), 5 μg/ml insulin (Sigma-Aldrich, I0516), and 1 μg/ml hydrocortisone (Sigma-Aldrich, H0888 – 1G). Cell lines were cultured at 37°C.

### RC-HiC library preparation

3-4×10^7^ number of 1-7HB2 cells were cross-linked on plate with formaldehyde (Agar Scientific R1026), followed by a quenching step with 1.25M glycine, scraping for cell detachment and two washes with cold PBS 1X. The cell pellet was re-suspended in 50ml freshly prepared ice-cold lysis buffer (10mM Tris-HCl pH 8, 10mM NaCl, 0.2% Igepal CA-630 (Sigma-Aldrich, I8896-50ML), one protease inhibitor cocktail tablet (Roche complete, EDTA-free 11873580001). Cells were lysed on ice for a total of 30min, with 2 x 10 strokes of a Dounce homogeniser with a 5-min break between Douncing to minimise cell clumping. Following lysis, the nuclei were pelleted and washed with cold 1.25xNEB Buffer 2 (NEB, B7002S) then re-suspended in 1.25xNEB Buffer 2 to make two aliquots of 10–15×10^6^ cells for digestion. Hi-C libraries were digested using 1500U MBOI (NEB, R0147M) at 37°C overnight while orbital shaking. Following digestion, the restriction fragment overhangs are filled in for 1h with dNTPs including biotin-14-dATP (Life Technologies, 19524-016). Fragments are then blunt-end ligated with 1U/μl T4 DNA ligase (Invitrogen, 15224-025) under dilute conditions to favour ligation between crosslinked fragments (in 15 ml tube for overnight at 16°C). DNA crosslinks were then reversed with 10mg/ml proteinase K (Roche, 03115879001) for 6-8 hours at 65°C followed by RNase A (Roche, 10109142001) treatment at 37°C for 60 minutes. Two rounds of DNA extraction/purification were carried out with phenol pH 8.0 (Sigma-Aldrich, P4557) and phenol: chloroform: isoamyl alcohol (Sigma-Aldrich, P2069) followed by precipitation with 3M sodium acetate pH 5.2 (Sigma-Aldrich, S7899) and 2.5x volume of ice cold 100% ethanol on wet ice for 1-2 hours. The Hi-C Library’s quantity and quality were assessed by running 50-100ng of the Hi-C libraries on a 0.8% agarose gel. Hi-C marking and Hi-C ligation efficiency was verified by PCR digest assay using MBOI and CLAI (NEB, R0197S) enzymes. Biotin is then removed from the ends of un-ligated fragments using the exonuclease properties of T4 DNA polymerase (NEB, M0203L) and DNA was sheared to obtain DNA fragments with a peak concentration around 400 bp. DNA ends were repaired and fragments, with internally incorporated biotin, are pulled down using magnetic Dynabeads MyOne Streptavidin T1 beads (Life Technologies 65601). After PE adaptors ligation ((5’-P-GATCGGAAGAGCGGTTCAGCAGGAATGCCGAG-3’ and 5’-ACACTCTTTCCCTACACGACGCTCTTCCGATC*T-3), pre-Capture amplification was performed with eight cycles of PCR on multiple parallel reactions from Hi-C libraries immobilized on Streptavidin beads, which were pooled post PCR and SPRI Ampure XP beads (Beckman Coulter, A63881) purified. The final Hi-C library was re-suspended in 25ul of Tris low-EDTA and quantified by the Qubit™ dsDNA BR Assay Kit (ThermoFisher, Q32853). The size distribution of the library was assessed by Tapestation D1000 (Agilent). The HiC capture regions were enriched via hybridisation with biotin-RNA probes (Agilent Technologies). The capture regions of interest were then pulled-down with Dynabeads MyOne Streptavidin T1 beads (Life Technologies 65601) and purified using SPRI Ampure XP beads (Beckman Coulter, A63881). Finally, Hi-C Capture library was amplified and then sequenced with Illumina 50 bp paired end sequencing.

### Capture biotinylated RNA oligos design

Capture biotinylated 120-mer RNA oligos (25–65% GC, <3 unknown (N) bases) were designed to target either one or both sides of MBOI site and within 4-500bp as close as possible to the ends of the targeted restriction fragments using a custom genome-wide Perl script made available from the Babraham Institute and then submitted to the Agilent eArray software (Agilent) for manufacture.

### RNA extraction and qPCR

RNA was extracted from cell lines using TRI Reagent (Sigma-Aldrich) and 1ug of total RNA was converted to cDNA using QuantiTect Reverse Transcription Kit (Qiagen). Quantitative PCR (qPCR) was performed using a ¼ dilution of cDNA with SYBR Green PCR Master Mix (Thermo Fisher Scientific) on ABI Step One Plus (Applied Biosystems) and specific primers (Sigma-Aldrich) for target genes (see SI Table 6).

### Public HiC Data

Published Human HiC data was obtained for three cell lines - GM12878, IMR-90, and H1-hESC from the 4D Nucleome project (Rao *et al.,* 2014; Dekker *et al.,* 2017).

### Public Gene Lists

Allele specific gene expression data for IMR-90 and H1-hESC were obtained from the UCSD Human Reference Epigenome Mapping Project (Kundaje *et al*., 2015) via the ASMdb (Zhou *et al*., 2022). Allele specific expression data for GM12878 were obtained from previously published work (see SI Table 2) (Workman *et al*., 2019). Known imprinted genes were obtained from (https://www.geneimprint.com). All genes were associated with their Ensembl ID in Gencode v38 (GRCh37) (Frankish *et al*., 2019). Genes with ambiguous or unknown Ensembl mapping were excluded from the analysis.

### Other Public Datasets

Allele-specific methylation datasets were obtained from Encode and previously published work via the ASMdb (see SI Table 3) (ENCODE, 2012; Kacmarczyk *et al*., 2018) CTCF datasets for GM12878 (GSM749704), IMR-90 (GSM935404) and H1-hESC (GSM733672) were obtained from Encode (ENCODE, 2012). CTCF data for 1-7HB2 was unavailable and we have instead used MCF10-A (GSE98551)(Fritz *et al*., 2018). For hromatin state data was obtained from the Roadmap Epigenomics Project (Kundaje *et al*., 2015). This dataset represents a core 15-state chromatin state model, built using ChromHMM (v1.10.0) (Ernst and Kellis, 2012), based on 5 epigenetic marks (H3K4me3, H3K4me1, H3K36me3, H3K27me3, H3K9me) (see SI Table 4). Chromatin loop data, generated using Peakachu (Salameh *et al*., 2020), for each cell line, were downloaded from the 3D Genome Browser (Wang *et al*., 2018). The allele-specificity of the loop, indicated by loop colour, is indicative of the subtraction matrix value at that loop position.

### Data Processing

HiC data was processed using our in-house pipeline, HiCFlow. Read adapters were trimmed, using Cutadapt (v3.5) (Martin, 2011) and truncated using HiCUP (v0.7.4) (Wingett *et al*., 2015) to remove sequences overlapping putative ligation sites. Processed reads were mapped independently to the GRCh37/h19 reference assembly using Bowtie2 (v2.4.4)(Langmead and Salzberg, 2012). The GRCh37 reference assembly was chosen as we observed mappability issues at the *IGF2-H19* locus in GRCh38. Since this locus is an essential control, we have herein presented all results using the GRCh37/hg19 reference. Alignment files were re-merged to paired-end files using Samtools (v1.1.0)(Li *et al*., 2009). Reads were deduplicated and processed to raw contact matrices using HiCExplorer (v3.7.1) (Zufferey *et al*., 2018). Finally, contact matrices were corrected using the KR balancing algorithm (Knight and Ruiz, 2013).

For allele-specific analysis, a phased haplotype for IMR-90 and H1-hESC was generated from the raw HiC data. Briefly, variants were called using the GATK (v4.2.4.1) best practises pipeline (Van der Auwera *et al*., 2013). Haplotype assembly was then performed using HapCUT2 (v1.3.2) (Edge, Bafna and Bansal, 2017). For GM12878, a phased haplotype was obtained from the Platinum Genomes phased variant truthset (Eberle *et al*., 2017). Prior to alignment, the reference genome was masked at site of phased variants using BEDTools (v2.29.2) to avoid reference bias during mapping (Quinlan and Hall, 2010). Finally, allelic assignment of reads was performed using SNPsplit (v0.5.0) (Krueger and Andrews, 2016). Visualisations were created using pyGenomeTracks (v3.6) (Lopez-Delisle *et al*., 2021).

#### Explanation of Subtraction matrices

Visual comparison of allelic matrices (A1 vs. A2) was performed using “subtraction matrices”. Joint-normalisation of raw A1 and A2 matrices was first performed using HiCcompare (v1.6.0) (Stansfield *et al*., 2018). This provides implicit correction of between sample bias. Normalised matrices were then transformed using the ‘Observed / Expected’ method to correct for genomic distance and more effectively resolve changes in long-range interactions. Normalised counts were subtracted (A2 – A1) and the resulting subtraction matrices were denoised using a median filter (Scikit-Learn v1.7.3) (Pedregosa *et al*., 2011). This emphasises regions with consistent directional bias and which are more likely to represent signals of interest (SI Fig 1c).

#### Bioinformatic processing of RC-HiC

*Processing of RC-HiC was as described above but analysis was restricted to read pairs mapping within a single capture region. All other read pairs were filtered from the analysis*

#### Quality Control

Following sequencing, quality control of the raw FASTQ data sets was performed using FastQC (v0.11.9). Screening for potential sequence contamination was performed using FastQ Screen (v0.5.2) (Wingett and Andrews, 2018). Reproducibility of the RC-HiC data set was assessed using HiCRep (v1.10.0) (Yang *et al*., 2017). Correlation between biological replicates was high (0.97 - 0.98 at 5kb resolution). Replicates were therefore merged in downstream analyses to improve resolution for haplotype phasing and allele-specific analysis.

### Compartment Analysis

HiCFlow performs compartment analysis using CscoreTool (v1.1) (Zheng and Zheng, 2018). Compartment analysis was performed on the full HiC datasets for each cell line at a 20kb resolution. The sign of the Cscore was oriented such that positive and negative scores represented “A” and “B” compartments respectively. Results were intersected with chromatin state data such that negative scores were associated with heterochromatin and positive scores were associated with active transcription (“activeTSS”).

### Allele-specific TAD (ASTAD) Classification

TAD domain detection was performed, using OnTAD (v1.2) (An *et al*., 2019), on the full HIC dataset binned at 10kb resolution. The set of TADs were then used as a reference set to identify TADs with substantial differences in contact frequency between allele specific matrices. A1 and A2 matrices were first jointly normalised using the LOESS method described in HiCcompare. A median filter was then applied to remove spurious or noisy background interactions. Following this, the absolute sum of differences is calculated within the relevant TAD domain. For each TAD, a Z-score is calculated by comparing against the chromosome-wide background level of absolute differences for a domain of equivalent size. The methodology of ASTAD detection is illustrated in Fig 4a.

### Enrichment Analysis

To determine if ASTADs were enriched for particular genomic features, we performed enrichment analysis using LOLA (v1.22)(Sheffield and Bock, 2016). ASTAD enrichment was tested against a background set of all identified TAD domains in a given cell line. To facilitate cell line comparison, only autosomal regions were tested for enrichment. A full list of genomic features tested and enrichment status is available in SI Table 5.

### Overlap Analysis

Overlap analysis was performed to identify ASTADs that were conserved between cell lines. Conserved ASTADs were defined as sets of ASTAD intervals, between cell lines, with at least a 90% reciprocal overlap. A 10% difference in overlap allows for slight error in domain positioning due the loss of resolution during matrix binning. Given a bin size of 20kb, a 10% difference equates to a shift of approximately one bin length for a median size domain interval. Interval overlap was calculated using BedTools (v2.29.2).

### Randomisation Testing

In each of the following randomisation tests any TAD domain overlapping a region with non-normal copy number or overlapping an Encode Blacklist region were removed from the enrichment analysis.

#### Comparison of identical heterozygous variants in conserved ASTAD boundaries compared to conserved TAD boundaries

Conserved identical heterozygous variants were defined as any heterozygotic variant (SNPs or Indels) present in all three cell lines. The total number of observed conserved variants overlapping the set of conserved ASTAD boundaries was compared against a null distribution of repeat random samples (n = 10,000) of conserved TAD boundaries. A TAD boundary was defined as the outer-most 20kb of a TAD domain, equivalent to 1 bin size on the AS-HiC matrices. This analysis was repeated for the Allele Specific Methylation (ASM) data for each cell line (see SI Table 8).

#### Enrichment of Imprinted (or ASE) genes in ASTADs relative to all TAD domains

The total number of observed Imprinted genes overlapping ASTADs was compared against an expected distribution of randomly sampled genes. Genes were selected via randomly stratified sampling to match the sample size and distribution of Imprinting gene types. Stratification was used due to the imbalance in gene type distributions between the Imprinted / ASE gene sets and the total gene sets. Genes were excluded from the analysis if they did not overlap a TAD domain or if they overlapped a blacklisted region or a region with non-normal copy number. A total of 10,000 samples were taken per analysis to build a null distribution and to calculate a Z-score from the observed overlap. This analysis was repeated for each cell line and for ASE genes (see SI Table 9).

### Methylation Status at CpG Islands

To check methylation status within regions of interest, preprocessed whole genome bisulfite sequencing (WGBS) data, corresponding to CpG methylation in ENCODE3 bed bedMethyl format, were obtained from ENCODE (GM12878: ENCSR890UQO, IMR-90: ENCSR888FON, H1-hESC: ENCSR617FKV). Filtering was performed using the methylKit R package (Akalin *et al*., 2012). For each cell line, all CpGs overlapping a CpG island were selected. For each target CpG island, boxplot was generated for comparison among three cell lines using the ANOVA statistics.

## Data availability

The dataset(s) supporting the conclusions of this article is(are) available in the **[repository name]** repository, **[unique persistent identifier and hyperlink to dataset(s) in http:// format]**. Scripts used for downstream bioinformatics analysis are available at https://github.com/StephenRicher/AS-HiC-Analysis (DOI: 10.5281/zenodo.6510198). Further details of the HiCFlow workflow are provided below. Any other relevant data are available from the corresponding authors upon reasonable request.

- Project name: HiCFlow
- Project home page: https://github.com/StephenRicher/HiCFlow
- Archived version: 10.5281/zenodo.6510187
- Operating system(s): Unix-based operating systems
- Programming language: Snakemake (Python)
- Other requirements: Snakemake 7.3.1 or higher, Conda
- License: MIT License
- Any restrictions to use by non-academics: None

## Author contributions

A.M., L.H. conceived and obtained grant funding the project, and supervised the research. A.M., G.P., and S.R. designed the study, G.P. acquired the experimental data, S.R. carried out bioinformatic and statistical analysis. A.M., L.H., G.P., S.R., T.Y. analysed data. S.S. design, troubleshooting, and logistics. A.M., G.P and S.R. wrote the manuscript. All the authors have read the manuscript.

## Acknowledgements

This work has been supported by the Medical Research Council (MR/P000711/1 to A.M. and L.H.), the Leverhulme Trust (RPG-2020-327 to A.M.), and the EPSRC DTP studentship (2106811 to S.R.). We acknowledge Dr Cameron Osborne (King’s College London) and Dr Valeriya Malysheva (University of Antwerp, Belgium) for guidance on preparing RC-HiC libraries and for critically reading the manuscript, Dr Zongling JI (University of Manchester) for providing H1-hESC RNA samples, and Dr Simon Andrews (The Babraham Institute Cambridge) for supplying the script to generate Hi-C probes.

---

**SI Fig 1:**
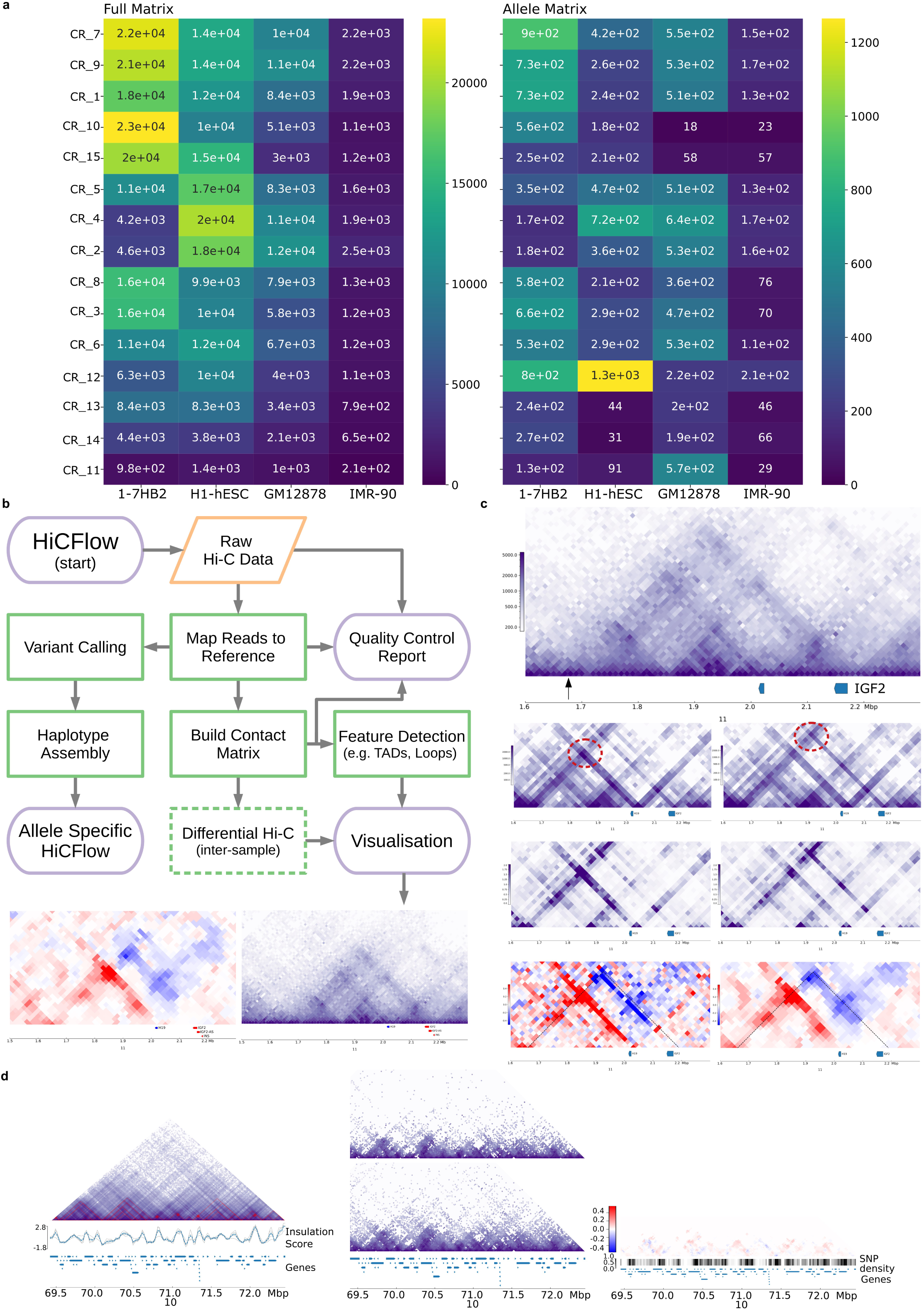
HiCFlow pipeline and Region Capture HiC (RC-HiC) library. **a)** Comparison of mean contact density (contacts/10kb) for RC-HiC (1-7HB2) across capture regions with the highest resolution human HiC datasets available: H1-hESC, GM12878 and IMR-90. In the full diploid matrix, the RC-HiC library yielded more contacts, at the given capture regions. The resolution of RC-HiC allele-specific matrices at the capture regions is comparable to GM12878, for which an experimental validated phased variant truth set is available. The capture region at chr6:26183471-26200857 did not yield sufficient data and was excluded. **b)** Overview of HiCFlow workflow – raw data is processed to publication ready visualisations or may be used for haplotype assembly in conjunction with allele-specific HiC analysis. See Methods for a full description of the bioinformatics tools used for each stage of the workflow. **c)** Illustration of method for generating subtraction matrix visualisations. The diploid HiC map (1) is split into two haploid matrices (2) using allelic-assignment via mate-rescue (Krueger and Andrews, 2016). Allele specific matrices (A1, A2) are normalised (3) by interaction distance using the method described in HiCExplorer – hicTransform (method=obs_exp) (Ramírez *et al*., 2018). Contact frequencies are subtracted (A2 – A1) (4) and the resulting matrices are de-noised (5), using a median filter (size = 3). This approach facilitates equal comparison of differences at all distances and removal of noisy interactions. **d)** A negative control region at chromosome 10, containing a high density of SNPs, showing that HiCFlow pipeline can split full matrix into haploid alleles (A1 and A2) but does not produce subtraction matrices where there is no allelic differences.

**SI Fig2:**
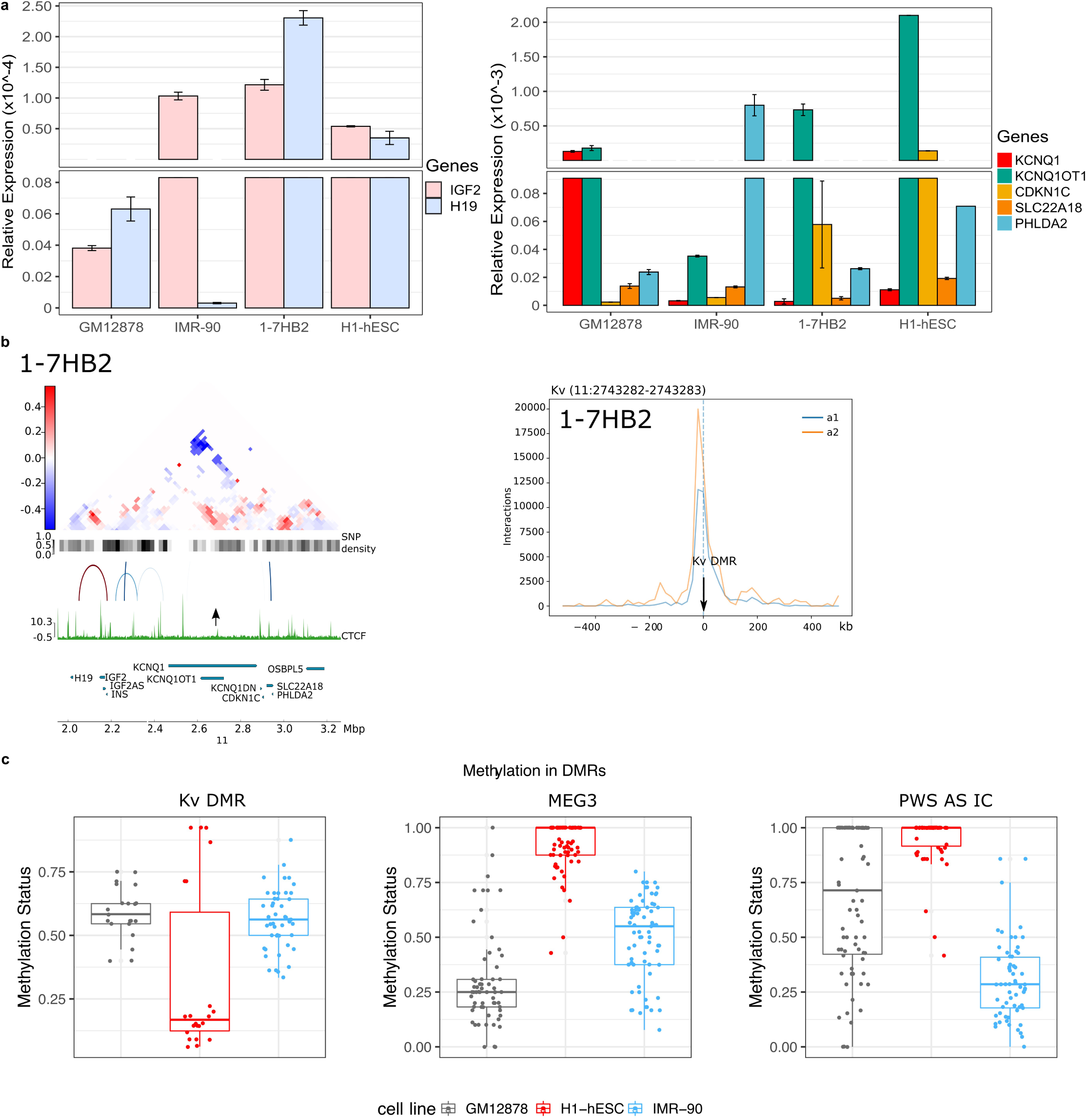
Supporting information relevant to Fig 2: DNA methylation data and expression levels of imprinted genes. **a)** Quantitative RT-PCR analysis of expression of imprinted genes inside of *H19-IGF2* and *KCNQ1* loci at the four cell lines considered in this study. **b**) 1-7HB2 data for KvDMR effect on allele specific associations showing subtraction matrix between A1 and A2 alleles and viewpoint analysis **c)** Full methylation profiles for CpG islands overlapping the imprinting control DMRs.

**SI Fig3:**
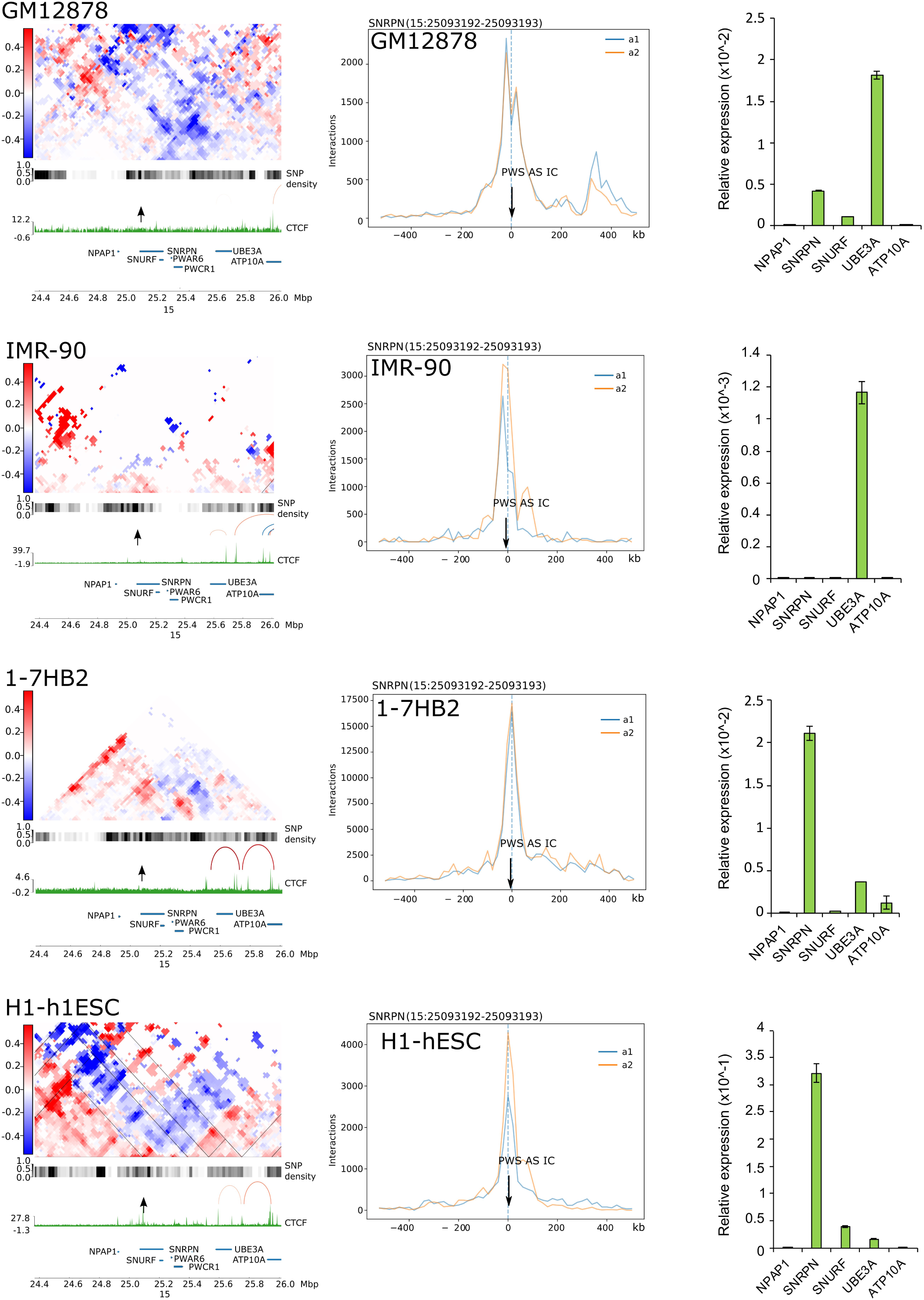
The effect of the PWS-AS imprinting control region on allele-specific chromatin conformation: Denoised subtraction matrix (left) and viewpoint analysis (middle) at the *SNRPN* locus, with underlying SNP density bars, CTCF tracks, allele-specific loops and imprinted gene tracks as in Fig 2. The ICR position is labelled with an arrow above the CTCF track, the coordinates are given above the viewpoint plots. Quantitative RT-PCR analysis of expression of imprinted genes at the *SNRPN* locus (right) in the four cell lines considered in this study.

**SI Fig4:**
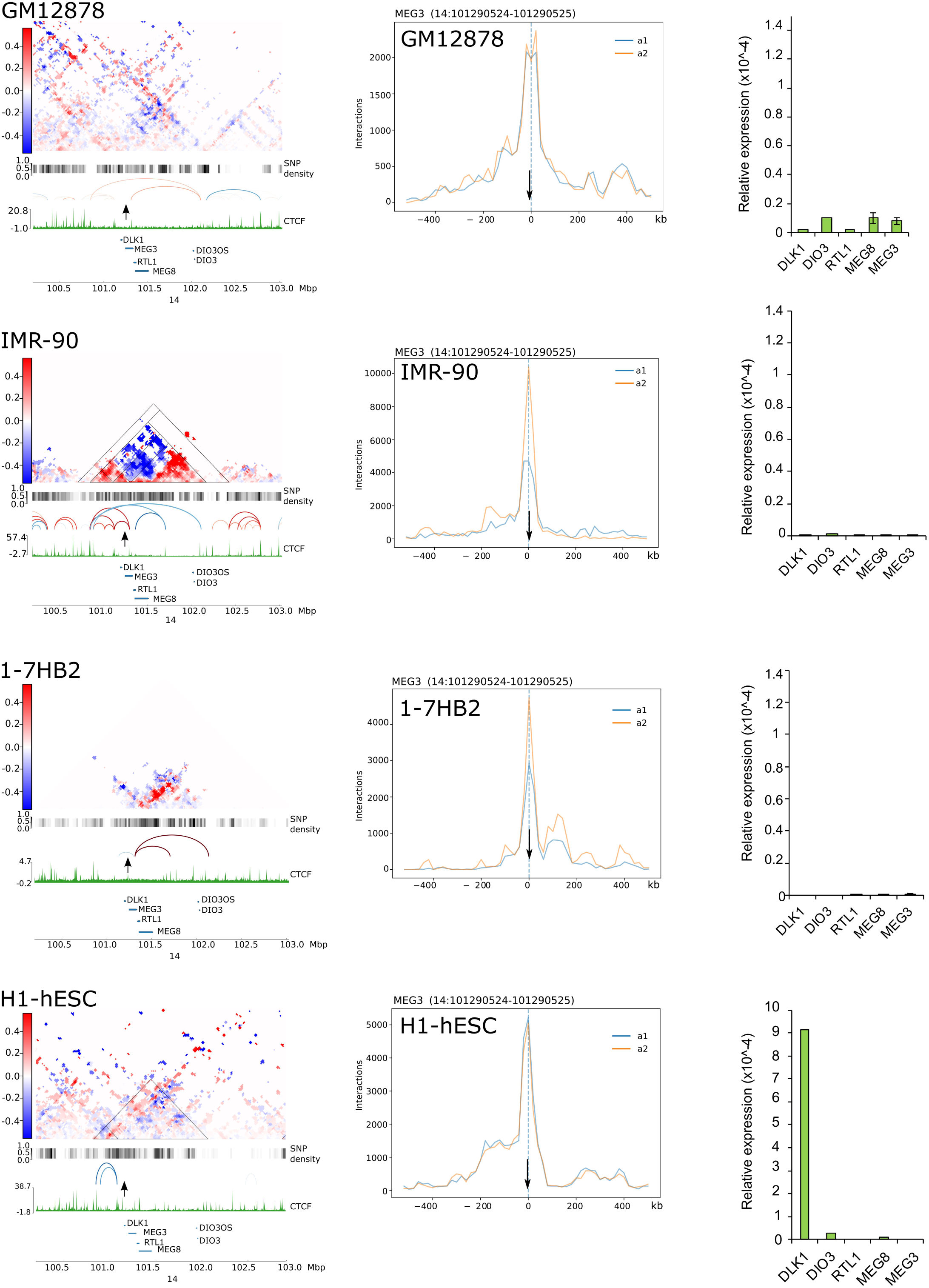
The effect of the IG-DMR/*MEG3* imprinting control region on allele-specific chromatin conformation: Denoised subtraction matrix (left) and viewpoint analysis (middle) at the *DLK1-DIO3* locus, with underlying SNP density bars, CTCF tracks, allele-specific loops and imprinted gene tracks as in Fig 2. The ICR position is labelled with an arrow above the CTCF track, the coordinates are given above the viewpoint plots. Quantitative RT-PCR analysis of expression of imprinted genes at the *DLK1-DIO3* locus (right) in the four cell lines considered in this study.

**SI Fig5:**
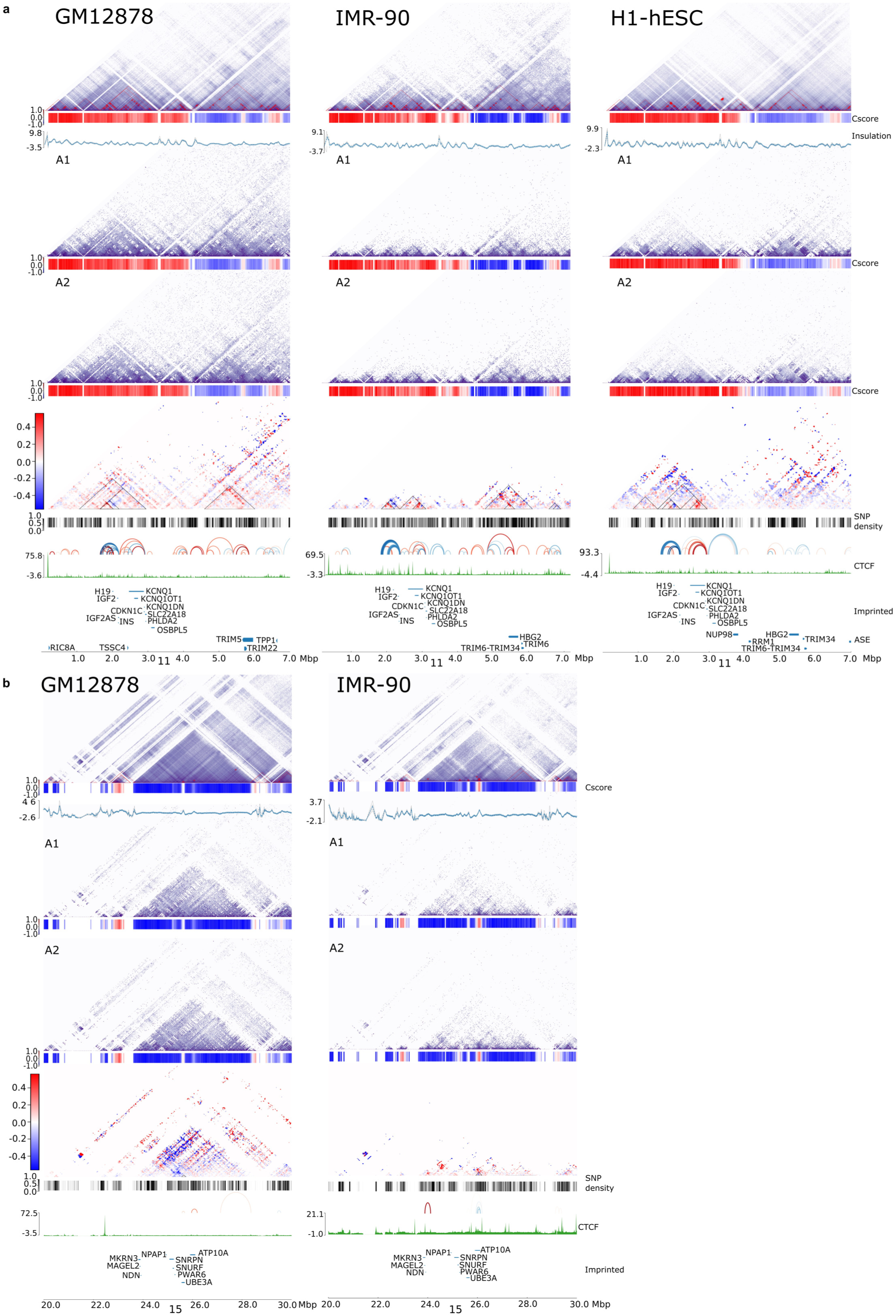
Compartment analysis of *H19-KCNQ1* and *SNRPN* loci: Full diploid and phased haploid (A1, A2) Hi-C matrices, with C-sores below indicating combined and allele-specific A-compartments (red) and B-compartments (blue) for wider regions around the imprinted domains. Subtraction matrices showing overall allelic differences, with SNP density, allele specific loops, CTCF tracks, imprinted gene positions as well as an additional track showing genes reported to have allele-specific expression in these cell lines. **a)** *H19-KCNQ1*, **b)** *SNRPN*.

**SI Fig6:**
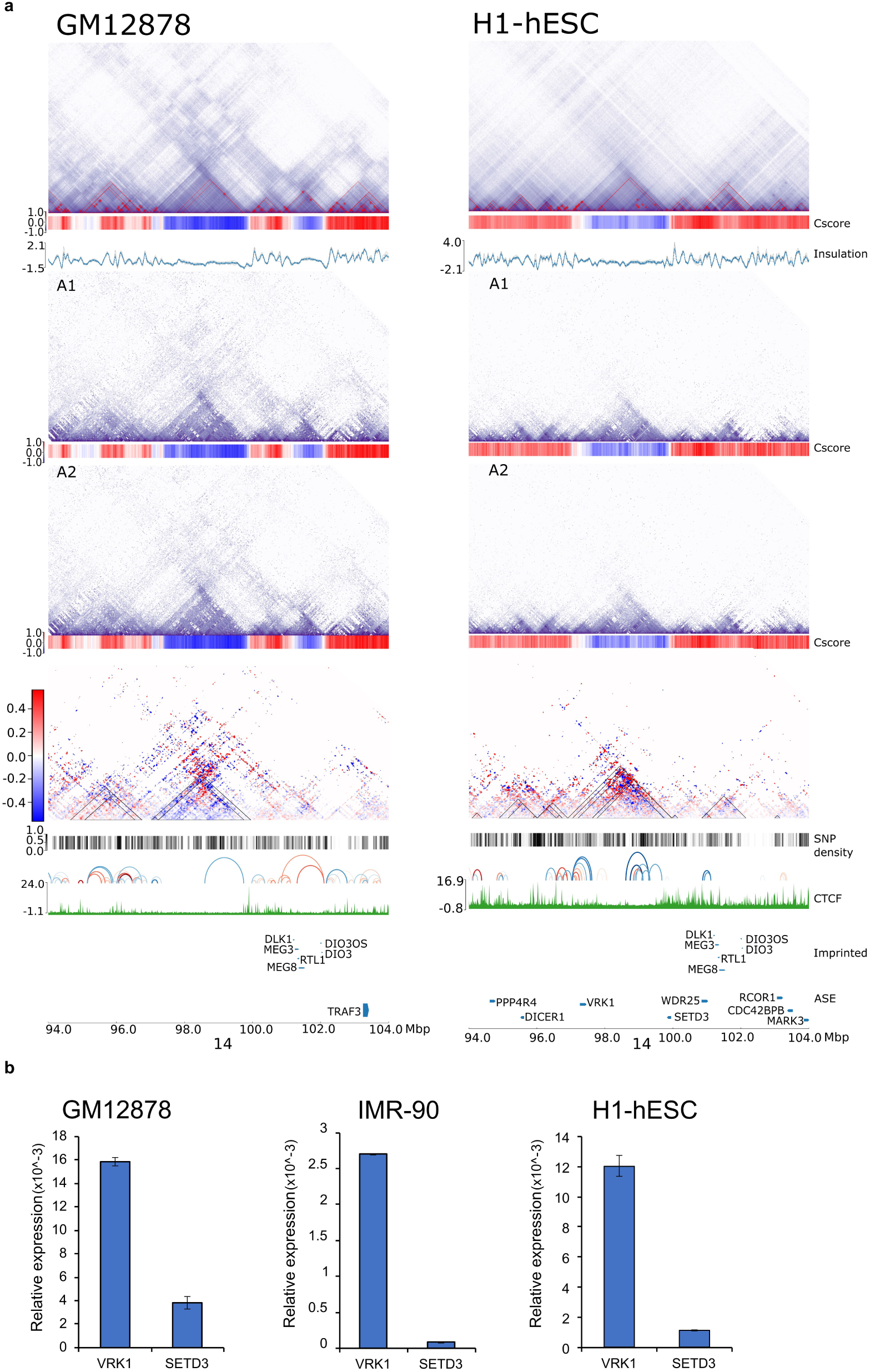
Compartment analysis of *DLK1-DIO3 locus*: Full diploid and phased haploid (A1, A2) Hi-C matrices, with C-sores below indicating combined and allele-specific A-compartments (red) and B-compartments (blue) for wider regions around the imprinted domains. Subtraction matrices showing overall allelic differences, with SNP density, allele specific loops, CTCF tracks, imprinted gene positions as well as an additional track showing genes reported to have allele-specific expression in these cell lines. **a)** *DLK1-DIO3* (right) loci in GM12878, IMR-90 and H1-hESC. **b)** Quantitative RT-PCR analysis of expression of *VRK1* and *SETD3* genes in GM12878, IMR-90 and H1-hESC cells.

**SI Fig7:**
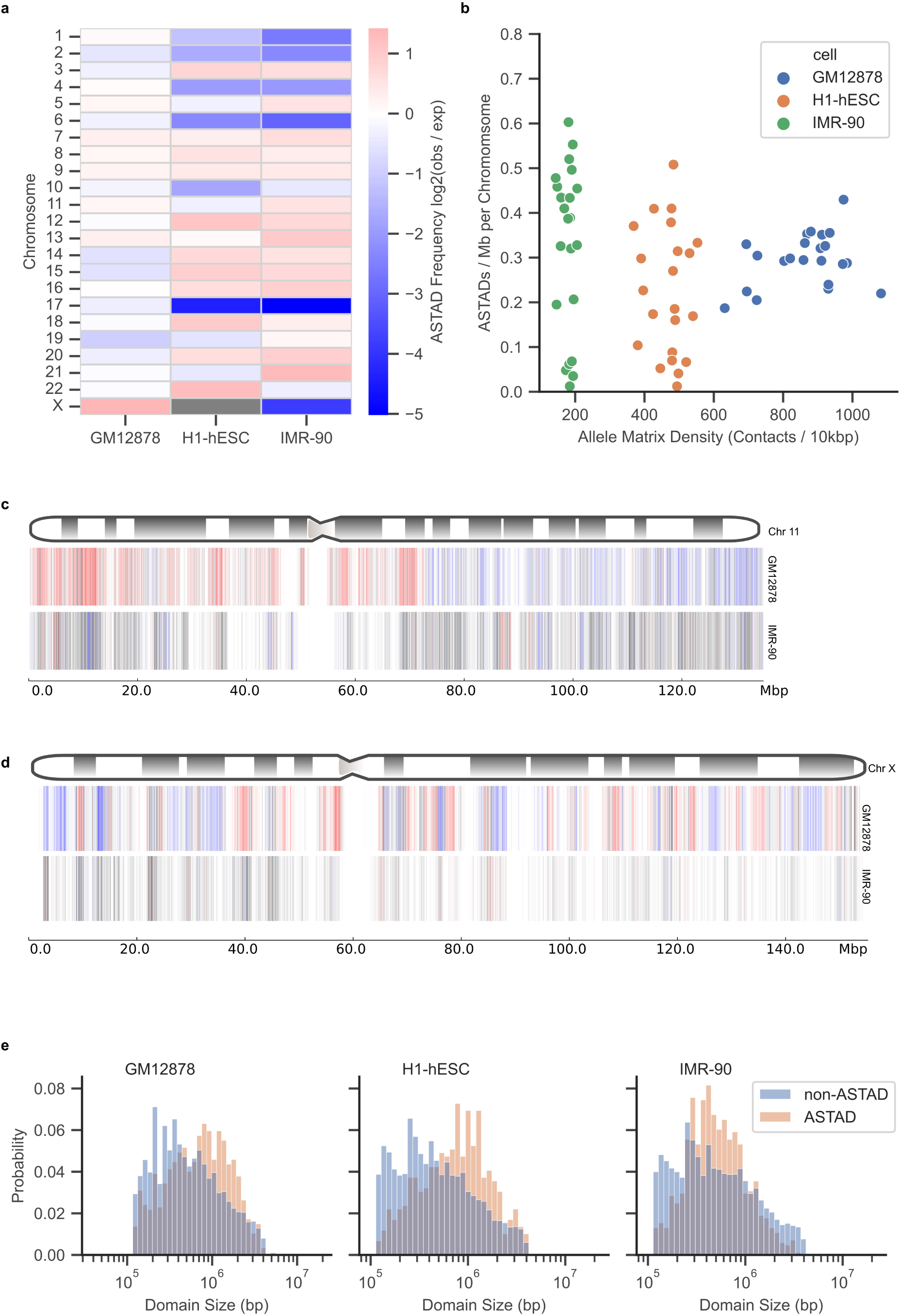
Features and distribution of ASTADs: **a)** ASTAD frequency relative to expected TAD proportions per chromosome. In GM12878, autosomal ASTAD distribution is similar to TAD distribution. The X-chromosome has a higher than expected proportion of ASTADs, likely due to skewed X-inactivation. In IMR-90 and H1-hESC ASTAD distribution is more variable with lower than expected proportions of ASTADs observed in chromosomes 1, 2, 4, 6 and 17. Lower than expected ASTADs on chromosome X in IMR-90 is consistent with random X-inactivation. **b)** Allelic contact frequency per-chromosome between cell lines. **c)** Ideogram of chromosome 11, with per-bin allelic differences for GM12878 and IMR-90. GM12878 chr11 is involved in a chromosome translocation leading to duplication of part of the q-arm. The ASTAD (A1-A2) change score is high along the entire region in GM12878, while in IMR-90 far fewer regions are highlighted. Regions with copy number changes have been excluded from ASTAD enrichment analysis. **d)** Ideogram of chromosome X with per-bin allelic differences for GM12878 and IMR-90 cells. GM12878 has non-random X-inactivation and shows substantial allelic differences along the chromosome. IMR-90, has random X-inactivation and HiCFlow does not the detect allele-specific differences in a whole cell population. **e)** Histogram of TAD / ASTAD domain sizes. ASTADs domains sizes are comparable to TADs, although there are lower proportions of smaller ASTADs.

**SI Fig8:**
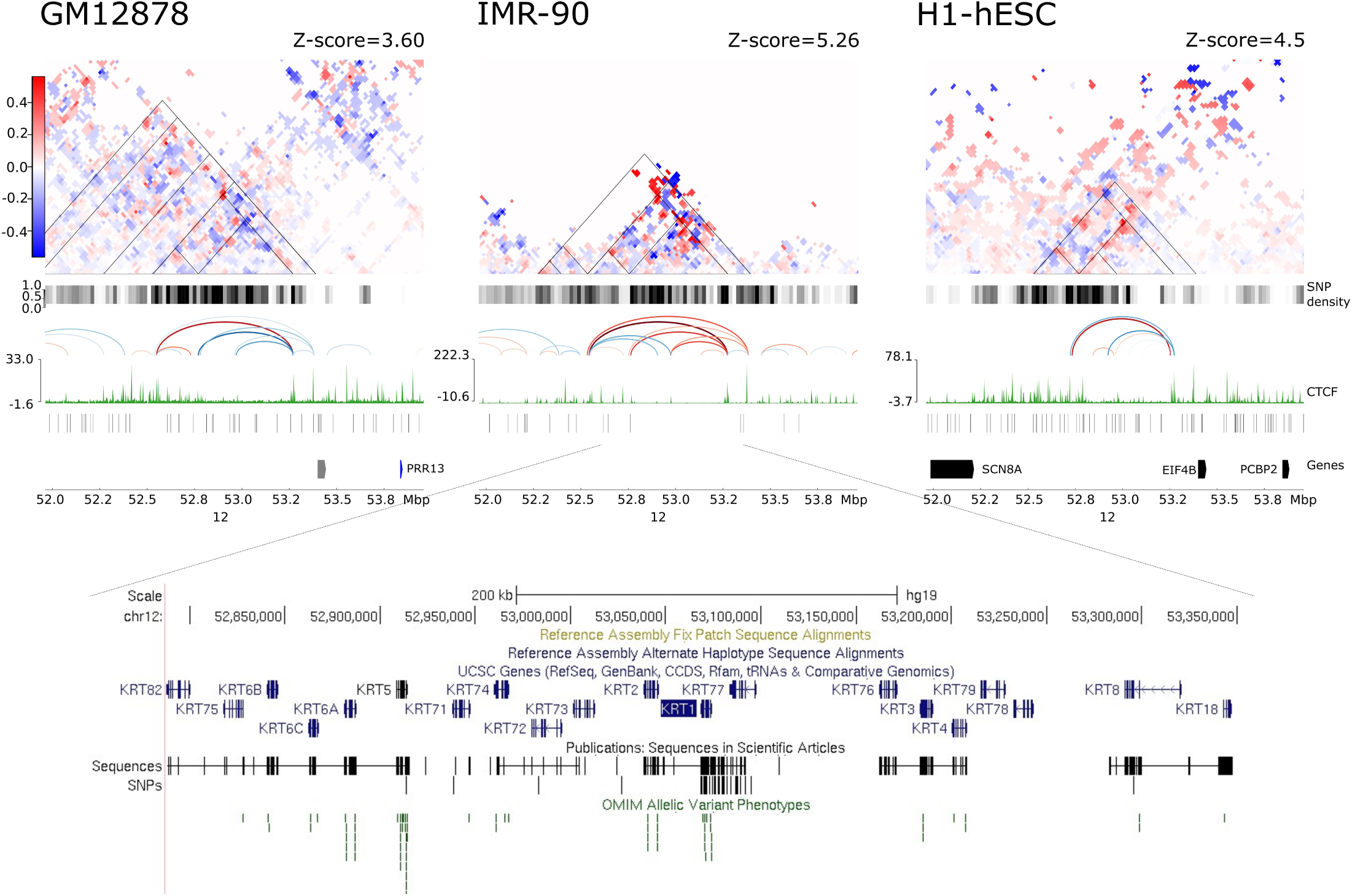
*KRT* gene cluster on chr12 is within a conserved ASTAD. Denoised subtraction matrices for the three cell lines showing ASTADs outlined as triangles. The expanded gene panel shows positions of the various *KRT* genes and density of OMIM allelic variant phenotypes within the conserved ASTAD (see SI File 5 for conserved variants at the ASTAD boundaries). Coordinates refer to genome build GRCh37/hg19.

**SI Table 1:**
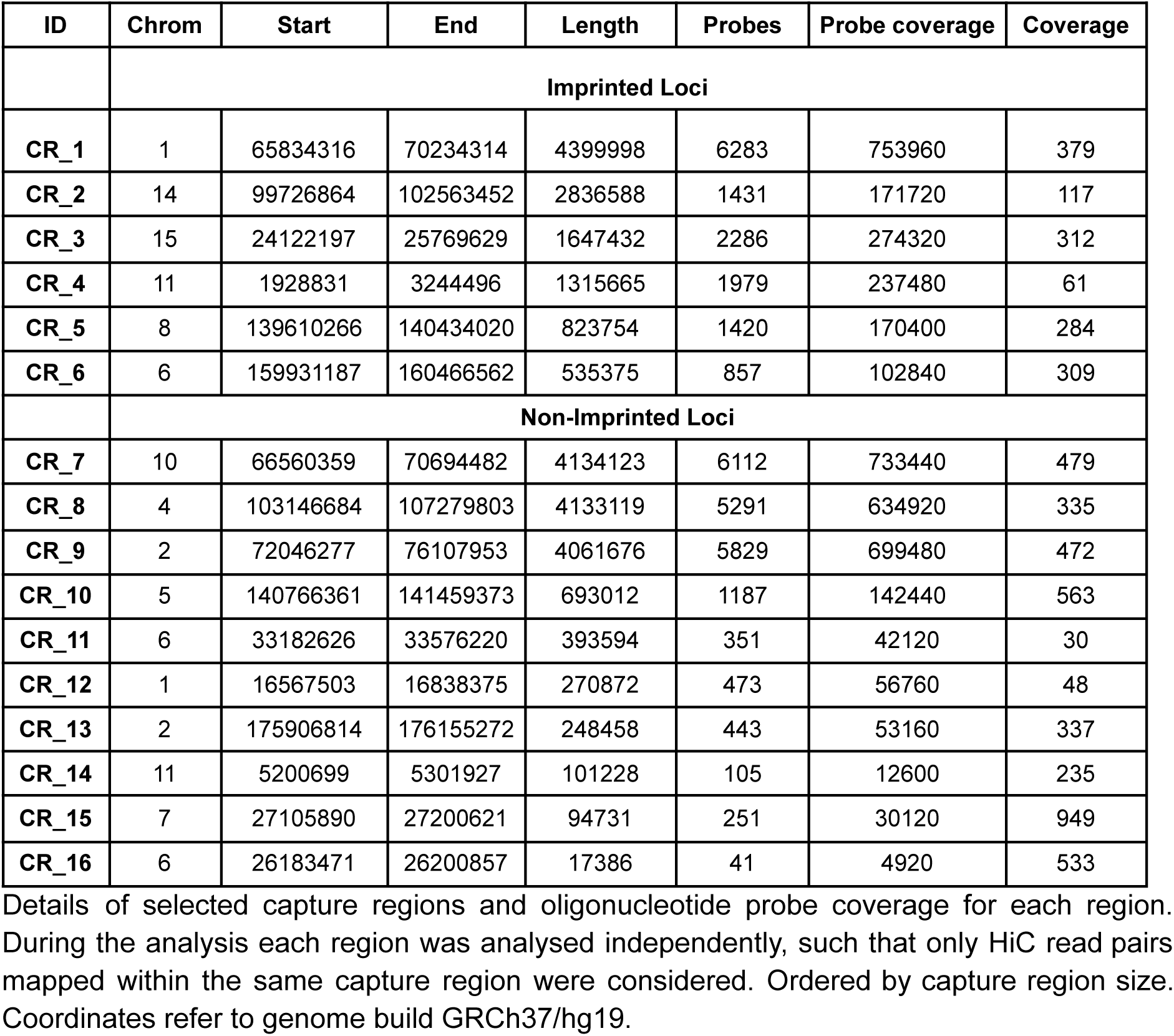
Capture Regions for RC-HiC in 1_7HB2 Cells.

**SI Table 2:** Allele Specific Gene Expression Data.

| Dataset | Total | Retained | Source | SNP Source |
| --- | --- | --- | --- | --- |
| **GM12878** | 482 | 480 | PMID:<br>31740818 | NA |
| **IMR-90** | 455 | 409 | GSM438363 | GSM432687 |
| **H1-hESC** | 2703 | 2398 | GSM438361 | GSM432685 |

Note: Allele specific gene expression (ASEG) data for GM12878 was obtained from previously published work as a list of 3592 Ensembl gene IDs which were found to contain at least two haplotype informative variants (Workman et al., 2019). Genes were filtered to retain only those genes (482) with allele specific expression bias (either paternal or maternal). The filtered gene list was merged with Ensembl IDs from Gencode v38. ASEG data for IMR90 and H1-hESC were obtained from the UCSD Human Reference Epigenome Mapping Project (Kundaje et al., 2015) via the ASMDb (Zhou et al., 2022). Genes were downloaded as a list of Gene symbols, which were mapped back to their associated Ensembl ID in Gencode v38. A number of genes had ambiguous or unknown Ensembl ID mappings and these were excluded from the final datasets.

**SI Table 3:** Allele Specific Methylation Data.

| Dataset | ASM Sites | ASM in CpG | Source | Source Size (Gbp) |
| --- | --- | --- | --- | --- |
| **GM12878** | 112910 | 865 | GSM1002650 | 120.4 |
| **IMR-90** | 13496 | 2355 | GSM2210597 | 70.7 |
| **H1-hESC** | 126023 | 656 | GSM1002649 | 127.9 |

**SI Table 4:** Core 15-state model (5 marks)

| State | Mnemonic | Description |
| --- | --- | --- |
| **1** | TssA | Active TSS |
| **2** | TssAFlnk | Flanking Active TSS |
| **3** | TxFlnk | Transcription at gene 5' and 3' |
| **4** | Tx | Strong transcription |
| **5** | TxWk | Weak transcription |
| **6** | EnhG | Genic enhancers |
| **7** | Enh | Enhancers |
| **8** | ZNF/Rpts | ZNF genes & repeats |
| **9** | Het | Heterochromatin |
| **10** | TssBiv | Bivalent / Poised TSS |
| **11** | BivFlnk | Flanking Bivalent TSS / Enh |
| **12** | EnhBiv | Bivalent Enhancer |
| **13** | ReprPC | Repressed PolyComb |
| **14** | ReprPCWk | Weak Repressed PolyComb |
| **15** | Quies | Quiescent / Low |

Note: Data was obtained, for each cell line, from the Roadmap Epigenomics Project (REF: Kundaje2015). 15 states were imputed based on 5 epigenetic marks using chromHMM (REF: Ernst2012).

**SI Table 5:** LOLA Enrichment “Max Rank” Score among 20 genomic features.

| Feature | GM12878 | H1-hESC | IMR-90 |
| --- | --- | --- | --- |
| **A Compartment** | **4** | **9** | **3** |
| **Weak Transcription** | **7** | 13 | **3** |
| **Weak Repressed Polycomb** | 11 | **10** | 14 |
| **CpG Islands** | 12 | 9 | 15 |
| **lncRNA** | 15 | **6** | 11 |
| **Flanking Active TSS** | **5** | 16 | **8** |
| **Repressed Polycomb** | 17 | **12** | 16 |
| **Heterochromatin** | **8** | **5** | 17 |
| **Bivalent Enhancer** | 18 | **13** | **15** |
| **Enhancers** | **3** | 18 | **3** |
| **ZNF / Genes Repeats** | **14** | 18 | **16** |
| **Protein Coding** | 13 | 19 | 17 |
| **Normal (CNV)** | 18 | **8** | 19 |
| **Flanking Bivalent TSS** | 19 | **15** | 19 |
| **Strong Transcription** | **12** | 20 | **11** |
| **Genic Enhancer** | 15 | 20 | **14** |
| **Bivalent Poised TSS** | 21 | 16 | 22 |
| **Quiescent** | 22 | 14 | 21 |
| **Transcription at 5' - 3'** | 17 | 22 | **20** |
| **Active TSS** | **9** | 22 | 13 |
| **Deletion (CNV)** | 22 | 21 | 23 |
| **B Compartment** | 23 | 23 | 23 |
| **Gain (CNV)** | 23 | 23 | **18** |
| **Loss (CNV)** | 24 | 24 | 24 |

Note: Enrichment of genomic features in ASTADs relative to TADs using LOLA (REF). Feature labelled according to their max LOLA ranking. Significant enrichment highlighted bold (alpha = 0.01, FDR corrected).

**SI Table 6:**
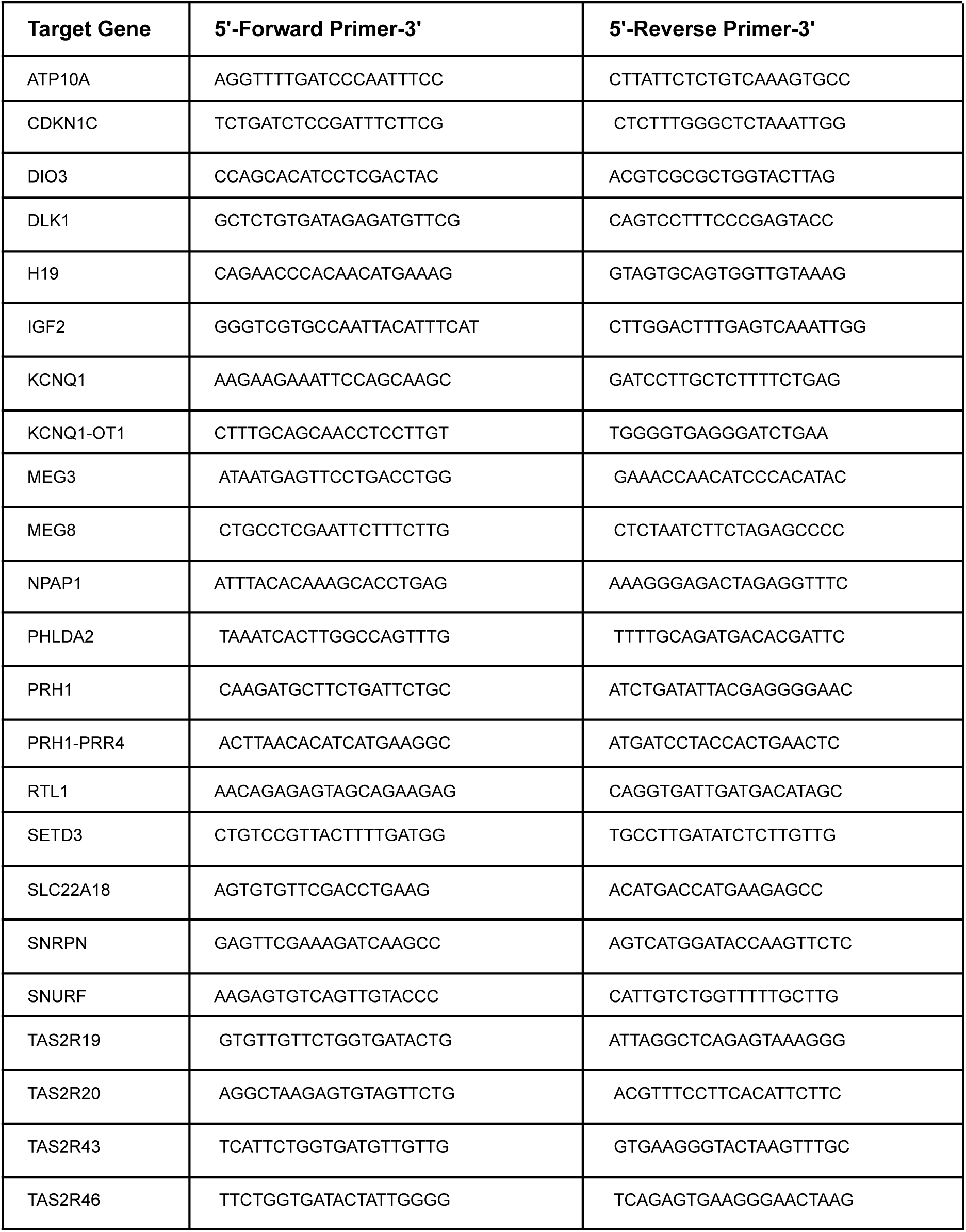
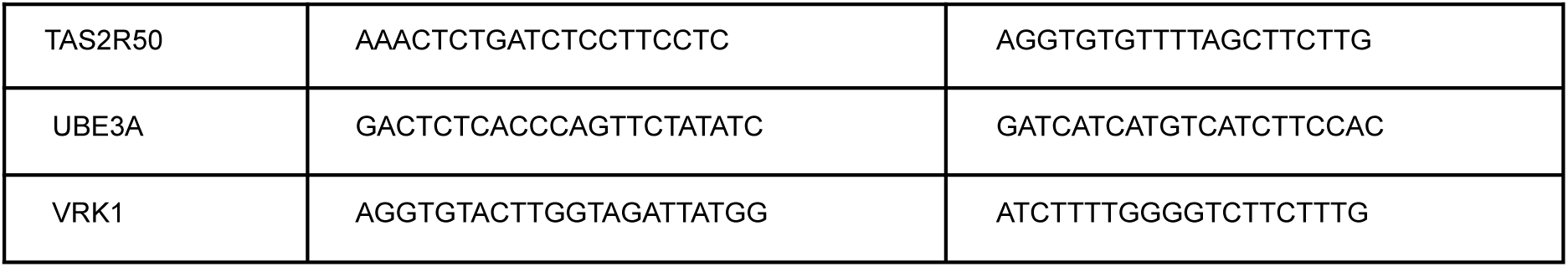
Oligo primers used in qPCR.

**SI Table 7:** Proportion of ASTAD / non-ASTAD domains overlapping Normal CNV.

|  | ASTAD (%) | All TADs (%) |
| --- | --- | --- |
| **GM12878** | 59 | 58 |
| **IMR-90** | 49 | 44 |
| **H1-hESC** | 76 | 72 |

Note: A domain was only considered overlapping if the entire span overlapped a region with Normal CNV. CNV detection was performed using the aligned HiC data by QDNAseq.

**SI Table 8:** Observed vs. Expected Overlap of Features in Conserved ASTADs.

| Test Name | Z score | P-value |
| --- | --- | --- |
| **Conserved Heterozygotic Variants** | 2.13 | 0.016 |
| **GM12878, Allele Specific Methylation** | 1.67 | 0.048 |
| **H1-hESC, Allele Specific Methylation** | -1.20 | 0.885 |
| **IMR-90, Allele Specific Methylation** | 1.27 | 0.102 |

Note: A domain was only considered overlapping if the entire span overlapped a region with Normal CNV. CNV detection was performed using the aligned HiC data by QDNAseq.

**SI Table 9:** Observed vs. Expected Overlap of Genes in ASTADs.

| Test Name | Z score | P-value |
| --- | --- | --- |
| **GM12878, Imprinted Genes** | 4.15 | < 0.001 |
| **GM12878, ASE Genes** | 4.59 | < 0.001 |
| **H1-hESC, Imprinted Genes** | 3.08 | 0.001 |
| **H1-hESC, ASE Genes** | -1.76 | 0.961 |
| **IMR-90, Imprinted Genes** | 5.00 | < 0.001 |
| **IMR-90, ASE Genes** | 3.50 | < 0.001 |

Note: The observed number of genes of interest (i.e. Imprinted) overlapping ASTADs was compared against an expected value determined by random sampling of all relevant genes (see methods). A null distribution of repeat samplings (n = 10,000) was built and a Z-score calculated to determine enrichment in ASTADs vs. TADs.

